# Correlation between circadian and photoperiodic latitudinal clines in *Drosophila littoralis*

**DOI:** 10.1101/2025.01.08.631867

**Authors:** Giulia Manoli, Pekka Lankinen, Enrico Bertolini, Charlotte Helfrich-Förster

## Abstract

Insects can survive harsh conditions, including Arctic winters, by entering a hormonally induced state of dormancy, known as diapause. Diapause is triggered by environmental cues such as shortening of the photoperiod (lengthening of the night). The time of entry into diapause depends on the latitude of the insects’ habitat, and this applies even within a species: populations living at higher latitudes enter diapause earlier in the year than populations living at lower latitudes. A long-standing question in biology is whether the internal circadian clock, which governs daily behaviour and serves as a reference clock to measure night length, shows similar latitudinal adaptations. To address this question, we examined the onset of diapause and various behavioural and molecular parameters of the circadian clock in the cosmopolitan fly, *Drosophila littoralis,* a species distributed throughout Europe from the Black Sea (41°N) to arctic regions (69°N). We found that all clock parameters examined showed the same correlation with latitude as the critical night length for diapause induction. We conclude that the circadian clock has adapted to the latitude and that this may result in the observed latitudinal differences in the onset of diapause.

## 1. Introduction

Insects have colonized virtually all ecosystems, including the Arctic regions, where they are exposed to cold and long winters. They hibernate in a state of dormancy (or diapause), which is hormonally induced and characterized by a reduced metabolism, increased stress resistance and an arrest in development or reproduction, enabling them to survive low temperatures and limited food supply (reviewed in [1]). It is important that insects prepare for diapause well before winter, as failure to adapt in time will ultimately lead to death. To do so they need a reliable indicator of the upcoming season, such as the shortening of the photoperiod (or lengthening of the night). The physiological responses of animals and plants to changes in day or night length are also known as photoperiodism. Many insects measure the length of the night and enter diapause when it exceeds a critical threshold – the critical night length (CNL) [2]. The CNL indicates the length of the night at which 50% of the insect population enters diapause. It strongly depends on latitude, since insects stemming from high latitudes need to enter diapause earlier in the year (at shorter CNL) than insects stemming from low latitudes [3–6]. So far, the physiological mechanisms underlying this latitudinal gradient in CNL are unknown.

A long-standing hypothesis is that the internal circadian clock, serves as time reference for measuring changes in night length through seasons [7]. If true, the circadian clock should also show differences between high- and low-latitude species. This is still being debated, since the circadian clock has other important functions, such as timing the daily activity-sleep cycle and many physiological parameters, and these should not vary too much at different latitudes [8]. Nevertheless, several studies reported latitudinal clines in clock properties such as robustness, period length and light sensitivity (reviewed by Hut et al. 2013 [9]). For example, several high-latitude *Drosophila* species possess weaker clocks than low-latitude species. This becomes evident in their locomotor activity or pupal eclosion rhythms that quickly become arrhythmic under constant conditions [10–15]. Furthermore, there are interesting differences in the anatomy and neurochemistry of the circadian clock neuronal network in high-latitude *Drosophila* species when compared to the low-latitude species *Drosophila melanogaster* [13,15–18]. For example, several high-latitude species virtually lack the neuropeptide Pigment-Dispersing Factor (PDF) in the main circadian pacemaker neurons, the s-LN_v_ (small ventrolateral neurons), which may explain the flies’ arrhythmic circadian behaviour [13,15–17,19]. PDF is also important to keep the low-latitude species *D. melanogaster* in the reproductive summer state by signalling to the neuroendocrine system (the insulin-producing median neurosecretory cells (MNCs)) [20–22]. The lack of PDF in the s-LN_v_ of high-latitude species may explain their strong diapausing phenotype. On the other hand, and unlike *D. melanogaster*, high-latitude flies express PDF in the corazonin (CRZ)-positive lateral neurosecretory cells (LNCs) [18]. Like the MNCs, the LNCs project to the *corpora cardiaca/allata* complex, where they release their neurohormones into the circulation. This makes the CRZ/PDF neurons interesting candidates for the regulation of the annual diapause in high-latitude flies.

Overall, there are interesting latitudinal differences in the circadian clock and its connections to the neuroendocrine system in different *Drosophila* species, which may explain the observed differences in their CNL for diapause induction. However, systematic studies of latitudinal clines in diapause and circadian clock organization in the same species are still lacking. We attempt to close this gap by studying the circadian clock properties of the fly *Drosophila littoralis*. *D. littoralis* belongs to the *virilis* group and is widely distributed throughout Europe, with the northernmost populations found in arctic regions (69°N) and the southernmost populations near the Black Sea (41°N, figure 1) [6,12]. This makes *D. littoralis* an ideal model for studying latitudinal clines. As other *Drosophila* species, *D. littoralis* overwinters as adult, and in females the arrest of reproduction during diapause is manifested by the atrophy of the ovaries [23,24]. *D. littoralis* strains from high latitudes enter diapause when the night lasts longer than 4 hours (corresponding to late summer), while strains from low latitudes do so when the night lasts longer than 12 hours (corresponding to fall) [6]. As true for other species of the *virilis* group, the circadian clock of *D. littoralis* seems to be weaker in the northern strains than in the southern ones as can be seen in their eclosion rhythms [12]. However, locomotor activity rhythms and the characteristics of the circadian clock in the brain have been studied only in one northern strain and not in other strains of this species [19].

**Figure 1.**
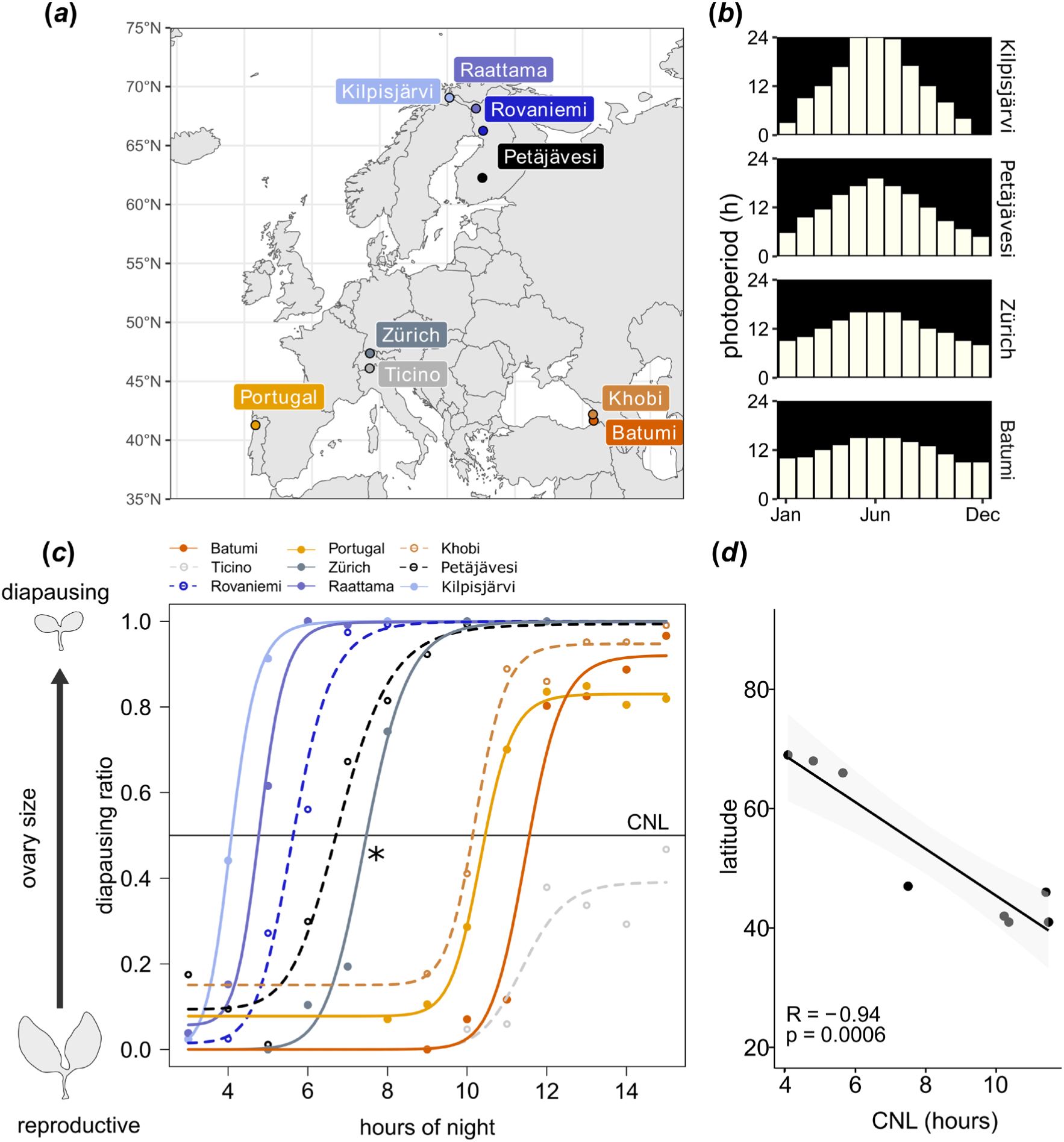
Photoperiodism of *D. littoralis* populations collected at different latitudes. (*a*) Latitudes at which the experimental lines were collected. (*b*) Mean photoperiod and night length by month at latitudes corresponding to some collection sites of the experimental lines. (*c*) Schematic of ovarian size under reproductive and diapausing conditions (left). The photoperiodic response curves of the different strains show latitude-dependent differences CNL (night length at which 50% of the population enters diapause) (right). The northern fly strains enter diapause at shorter CNLs than fly populations collected at central or southern latitudes, which have longer CNLs. An outlier (asterisk) is the strain Zürich that has an unusually short CNL for its rather southern location. This may be caused by the relatively high altitude (∼1000m) in the Alpes that causes lower temperatures. In contrast, the strain Ticino, which originates from a similar latitude as Zürich never reaches 100% diapause. This may be explained by the exceptionally mild climate of this region (∼200m altitude), where temperatures rarely fall below 0°C. *(d)* A Pearson correlation test revealed a significant negative correlation between latitude and CNL: CNL increased with decreasing latitudes. R: Correlation coefficient, p: Significance level.

In this study, we investigated whether the latitudinal differences in the onset of diapause are correlated with differences in the circadian clock of *D. littoralis*. Therefore, we analysed the locomotor activity rhythms and the molecular clock oscillations in adults of selected *D. littoralis* strains originating from latitudes between 41°N and 69°N. Furthermore, we analysed the circadian oscillations of PDF in the s-LN_v_ terminals of a southern and a northern *D. littoralis* strain and investigated whether the photoperiod affects the amount of PDF in these terminals as well as the amount of PDF and CRZ in the LNCs. We found a significant correlation between circadian parameters and the timing of diapause and have the first evidence that PDF in the CRZ neurons of northern *D. littoralis* may have taken over the function of PDF in the s-LN_v_ terminals of southern strains.

## 2. Results

### 2.1. Latitude-dependent variations in photoperiodic diapause of *D. littoralis*

We investigated the photoperiodic response of *D. littoralis* strains collected across a latitudinal gradient in Europe (figure 1*a*) by exposing flies to days with different photoperiods but a constant temperature of 16°C, thus mimicking summer-like (long days and short nights) or winter-like (short days and long nights) light conditions in the laboratory. Photoperiods that the flies experience in nature are shown for four strains stemming from different latitudes in figure 1*b*.

Photoperiodism in flies was assessed by evaluating the stage of ovarian maturation in females, which is a marker of reproductive diapause [24]. To determine the CNLs at which 50% of the flies of each population enter diapause, we generated photoperiodic response curves for six previously studied strains (Kilpisjärvi, Rovaniemi, Zürich, Ticino, Khobi and Batumi) [12] and three more recently collected strains (Portugal, Petäjävesi and Raattama) (figure 1*c*; table S1). Correlation analysis revealed a negative correlation between latitude of origin and CNL (figure 1*d*). This is consistent with previously reported data by Lumme and Oikarinen [6] and Lankinen [12], who found a latitudinal cline in the onset of diapause of *D. littoralis*. Importantly, the diapause levels of the previously studied strains still follow the same latitudinal cline as previously found [12], despite the long time in captivity, and the three new strains fit well into it, as can be seen from the photoperiodic response curves of all strains (figure 1*c*; table S1).

Overall, we have established a panel of laboratory inbred *D. littoralis* strains with a robust photoperiodic phenotype and a clear latitudinal cline in the CNL (figure 1*d*).

### 2.2. Latitude-dependent variations in circadian rhythmicity of *D. littoralis*

To test whether circadian behaviours of *D. littoralis* vary also with the latitude of origin, we recorded locomotor activity of five selected *D. littoralis* strains (Kilpisjärvi, Raattama, Zürich, Portugal, and Batumi) in the *Drosophila* Activity Monitor system (Trikinetics, Princeton, MA, USA) at three different photoperiods (LD8:16, LD12:12 or LD20:4). *D. melanogaster*, the common laboratory fruit fly, served as a reference as most of the knowledge about the circadian clock in flies derives from this species [25]. Remarkably, a photoperiod of LD12:12 (12 hours of light followed by 12 hours of darkness, equinox-like) corresponds to winter light conditions at high latitudes, whereas at low latitudes it corresponds to light conditions of spring and fall days. Furthermore, extreme photoperiods such as LD20:4 are typical summer photoperiods for the northern *D. littoralis* strains, but these light conditions are not experienced in nature by the southern European strains. In contrast, LD8:16 is a short photoperiod that occurs throughout continental Europe and in the northern part of the *D. littoralis* range, but that will not impact locomotion of northern populations because under this photoperiod flies are inactive (diapausing).

Individual flies were recorded for five days under LD8:16, LD12:12 or LD20:4 and then released into constant darkness (DD) to assess their free-running rhythmicity in the absence of external environmental cues. The rhythms of the different fly populations are shown in figure 2*a*, while actograms of individual flies are shown in figure S1. The Lomb-Scargle periodogram was used to quantify the endogenous period length (figure 2*b*, table S3) and to determine the percentage of rhythmic flies for each strain (figure 2*c*). As shown previously, *D. melanogaster* flies are rhythmic when released in DD, regardless of the photoperiod to which they were previously exposed [26]. In contrast, southern *D. littoralis* strains (Batumi and Portugal) showed a higher rhythmicity in DD when they had previously experienced short, but not long, photoperiods (figure 2*a-c*). The loss of rhythmicity after exposure to long photoperiods likely reflects the adaptation of the circadian clock of southern flies to only moderate changes in day lengths. *D. littoralis* strains from the northernmost collection sites (Kilpisjärvi and Raattama), however, rapidly became arrhythmic under DD after all three photoperiods. Behavioural arrhythmicity under constant conditions of northern fly populations is a known trait which has been reported for several species [13–16,19]. Here we show that within a single species this trait is plastic and latitude dependent. The correlation analysis revealed a significant negative correlation between circadian rhythmicity in DD and the latitudes from which the strains originated (figure 2*h*). The free-running periods of the rhythmic flies were not significantly different in the *D. littoralis* strains (table S3).

**Figure 2.**
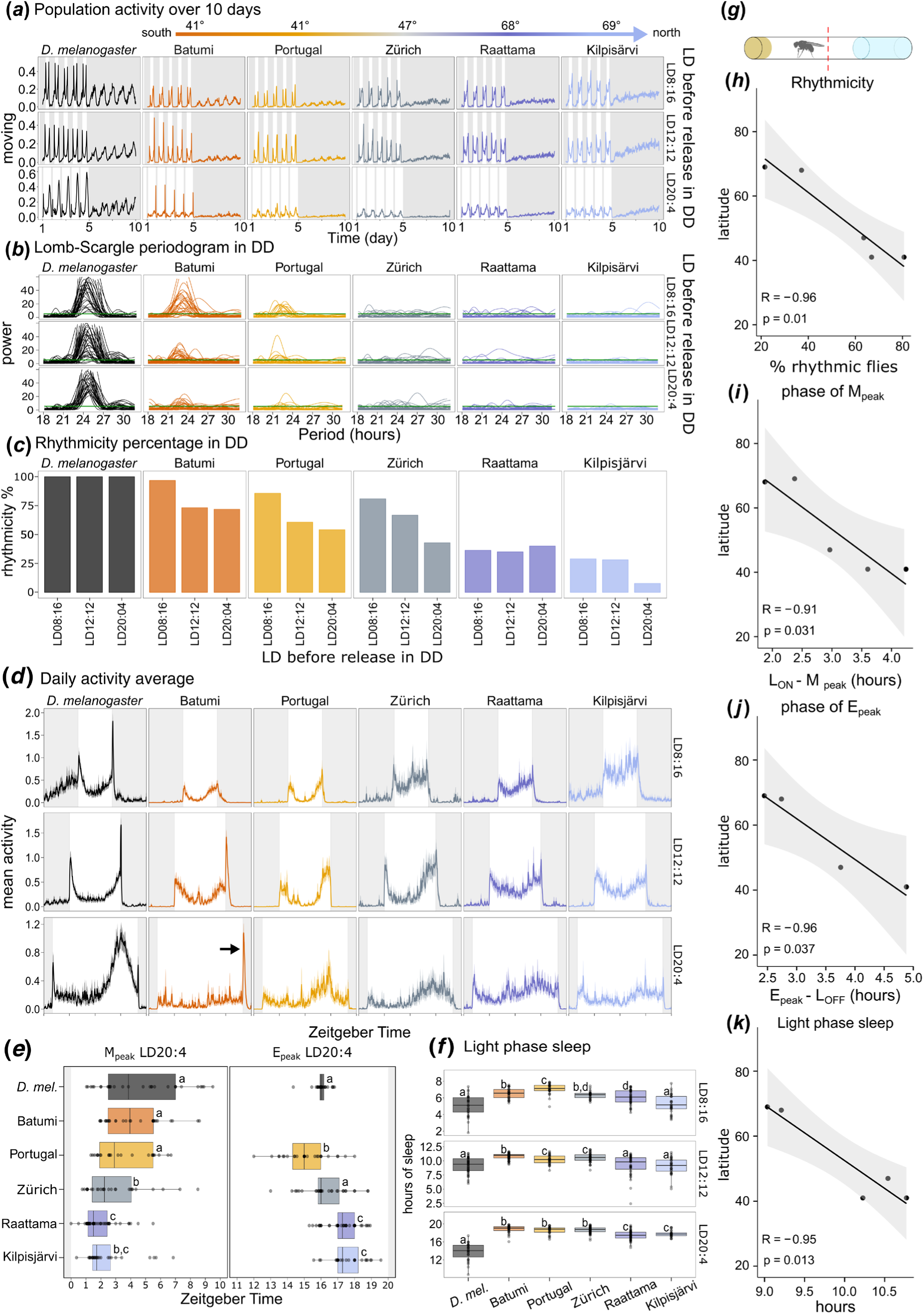
Circadian rhythmic parameters of *D. melanogaster* and *D. littoralis* from southern and northern latitudes exposed to different photoperiods and released in DD. (*a*) Average population activity of about 30 flies over 15 days; the first 5 days the flies were kept under LD cycles; from day 5 on they were released in DD (shaded area). The precise number of tested flies is indicated in table S1 and S2. The *D. melanogaster* population was always strongly rhythmic in DD, while the southern *D. littoralis* populations were less rhythmic after being exposed to LD20:4. The northern *D. littoralis* populations became quickly arrhythmic in DD. (*b*) Lomb-Scargle periodograms in DD, superimposed for all flies of a strain. (*c*) Percentage of rhythmic flies under DD. (*d*) Average daily activity profiles of all strains. For detailed explanations see the text. (*e*) Timing of M_peak_ and E_peak_ in LD20:4. M_peaks_ became earlier and E_peaks_ later with increasing latitude of the strains’ origin. Different letters indicate significant differences between the different strains (after Kruskal-Wallis p<0.05). The E_peak_ could not be determined for Batumi because of the prominent L_OFF_ peak (arrow in panel (*d)*). (*f*) Under every condition tested, the amount of light-phase sleep was higher in the southern strains (Batumi and Portugal) compared to the northern ones (Kilpisjärvi and Raattama), after Kruskal-Wallis p<0.05. (*g*) Schematic of the DAM recording system. A single fly is isolated in a tube with food in one end and a foam plug in the other. The locomotor activity is a count of the interruption of the infrared beam in the middle of the tube per minute. (*h*) Correlation between latitude and rhythmicity (average of the different conditions tested). (*i*) Correlation between latitude and the timing of morning activity in relation to light-on (L_ON_ - M_peak_) in LD20:4. (*j*) Correlation between latitude and the timing of evening activity in relation to light-off (E_peak_ - L_OFF_) in LD20:4. (*k*) Correlation between latitude and light-phase sleep. The Pearson correlation test showed a significant negative correlation for all parameters with latitude, meaning that there was an increase in the values with decreasing latitudes.

### 2.3. Latitude-dependent variations in daily activity patterns of *D. littoralis*

To extract further behavioural markers known to be under the control of the circadian clock, we analysed the average activity profiles of the different *D. littoralis* strains under the LD conditions that preceded their release into DD. We found that southern strains showed a bimodal activity pattern with a clear morning peak (M_peak_) and evening peak (E_peak_) and a reduced activity (increased sleep) in the middle of the light phase (siesta or light-phase sleep). The northern strains (Raattama and Kilpisjärvi) instead showed a higher variability in activity between individuals, resulting in noisier activity patterns (figure 2*d*). The strain from Zürich showed an intermediate activity profile between the southern and northern strains (figure 2*d*) as was already seen for its rhythmicity in DD (figure 2*a-c*). Notably, under LD8:16, the activity pattern of the southern *D. littoralis* strains (Batumi and Portugal) was more similar to the activity of *D. melanogaster* than to that of the northern *D. littoralis* strains (Raattama and Kilpisjärvi). Under LD12:12, all strains showed some degree of bimodal activity, with the northern *D. littoralis* strains showing higher activity levels with less pronounced siesta (figure 2*d, f*). At LD20:4, the siesta (light-phase sleep) became less pronounced in all flies, but it remained less deep in the northern strains (figures 2d, *f*, S2), revealing a clear latitudinal cline (figure 2*k*). In addition, major differences between *D. melanogaster* and the different *D. littoralis* strains occurred in the timing of M_peak_ and E_peak_ under LD20:4. *D. melanogaster* and the southern *D. littoralis* strains from Batumi and Portugal showed only little morning activity, which, if present, started well after light-on (L_ON_). Evening activity, on the other hand, started early and decreased already before light-off (L_OFF_). The northern strains (Raattama and Kilpisjärvi) showed instead a more pronounced M activity shortly after L_ON_ and an E activity that remained high until the end of the day (figure 2*d*). The quantification of the M_peak_ and E_peak_ under the longest photoperiod (LD20:4) showed a late M_peak_ and early E_peak_ in the southern strain (Portugal), an intermediate phenotype in the central Europe strain (Zürich) but an early M_peak_ and late E_peak_ in the northern ones (Raattama and Kilpisjärvi; figure 2*e*). For the southern Batumi strain, we could not determine the E_peak_ because it was masked by the strong L_OFF_ peak (see below). The timing of M_peak_ and E_peak_ of the northern strains was more plastic than that of the southern strains meaning that they could better follow L_ON_ and L_OFF_ under very long photoperiods. The values of the estimated M and E peaks were significantly different between northern and southern *D. littoralis* strain tested and showed a clear latitudinal cline (figure 2*i, j*).

Other features of the daily activity patterns of *Drosophila* such as the pronounced sudden increases of locomotor activity directly after L_ON_ and L_OFF_ transitions are known to be independent of the circadian clock [27,28]. In *D. melanogaster* these L_ON_ and L_OFF_ peaks, also referred to as startle responses, were present under every photoperiod tested (figure 2*d*).

Interestingly, the startle response at L_ON_ is completely absent in all *D. littoralis* strains, while the startle response at L_OFF_ is reduced in most strains except for the southern Batumi strain (figure 2*d*). Absence of the startle responses has already been reported in other fly species of the *virilis* group collected at high latitudes, such as *Drosophila lummei* and *D. ezoana* [11,19], suggesting that this is due to a decrease in light sensitivity with increasing latitude.

In summary, we found a negative correlation of several clock-dependent phenotypes with the latitude of origin, such as the average percentage of rhythmicity in DD (figure 2*h*), phase of the M_peak_ (figure 2*i*) and the E_peak_ (figure 2*j*), and sleep hours during the light phase (figure 2*k*). All these measured values decreased with increasing latitude. The present results illustrate that the adaptation of *D. littoralis* to different latitudes is not only reflected in seasonal responses such as reproductive diapause (figure 1), but also in the circadian clock that controls daily behaviour (figure 2).

### 2.4. Latitude-dependent variations in clock protein cycling of D*. littoralis*

We previously found that in the northern *D. littoralis* strain Kilpisjärvi the circadian clock protein PERIOD (PER) was expressed in all circadian clock neurons known in *D. melanogaster* (figure 3*a*), with the exception of the dorsal clock neurons 2 (DN_2_) and the large ventrolateral neurons (l-LN_v_) [18]. Here, we extended this analysis to a southern *D. littoralis* strain (Batumi) and additionally performed immunostaining against the clock components Par Domain Protein 1 (PDP1) and Pigment-Dispersing Factor (PDF). In the DN_2_ of both *D. littoralis* strains, we still could not detect clear staining, but PDP1 and PDF staining were present in the l-LN_v_ of both strains (figure S3), suggesting that all clock neurons are present in *D. littoralis*, but that some of them express only small amounts of clock proteins and are therefore difficult to detect.

**Figure 3.**
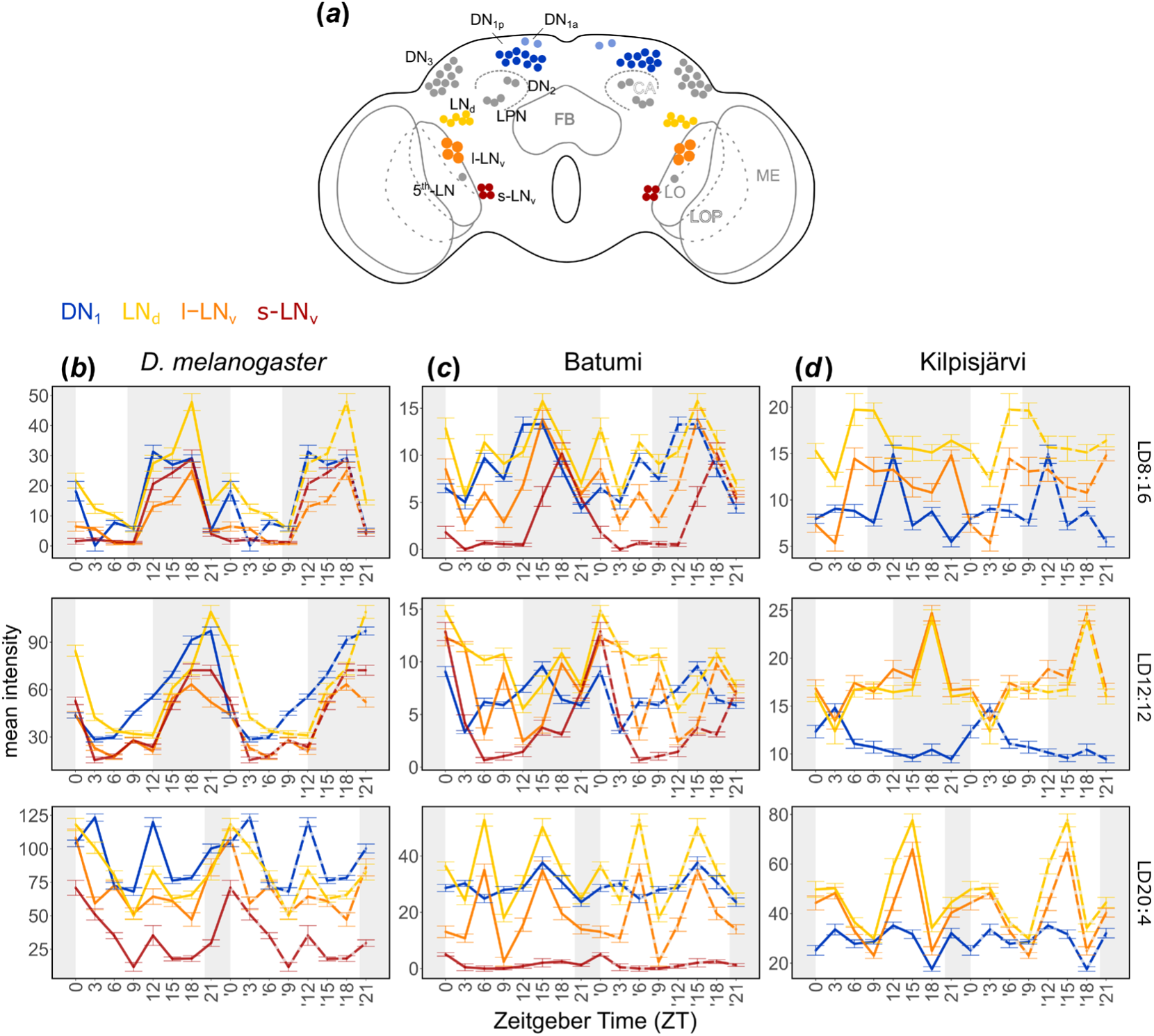
PDP1 cycling in different clock neuronal clusters of *D. melanogaster, D. littoralis* north (Kilpisjärvi) and south (Batumi) under different photoperiods. The staining values were double-plotted (dashed lines) to improve visualization. (*a*) Clock neurons of *Drosophila*. (b) PDP1 oscillations in the brain of *D. melanogaster* (average of 10 brains). Under LD8:16 and LD12:12 the different clock neurons showed an overall synchronized PDP1 cycling. Under the longest photoperiod LD20:4, the synchrony is partially lost and the cycle of PDP1 looked bimodal. (*c*) PDP1 oscillations in the brain of *D. littoralis* from Batumi (average of 10 brains). Under LD8:16 most of the neurons are synchronized in the PDP1 cycle. Under LD12:12 the synchrony was still present but to a lower degree and it was strongly reduced under LD20:4. (*d*) PDP1 oscillations in the brain of *D. littoralis* from Kilpisjärvi (average of 10 brains). PDP1 was absent in the s-LN_v_.Under LD8:16 and LD12:12 the PDP1 cycle in the other neurons was poorly synchronized, however it became more synchronous under the longest photoperiod (LD20:4).

We performed time series of immunostaining in *D. littoralis* from Kilpisjärvi and Batumi, with an antibody against PDP1 under three photoperiods (LD8:16, LD12:12 and LD20:4) to quantify circadian protein oscillation in the clock neurons. PDP1 staining was quantified in three groups of lateral clock neurons (the l-LN_v_, s-LN_v_ and the dorsolateral neurons, LN_d_) as these groups are responsible for timing M and E activity peaks in *D. melanogaster* [29], and PDP1 has been shown to oscillate strongly in them [30]. Furthermore, we quantified PDP1 staining in one group of dorsal neurons, the DN_1_, which is of similar importance in the control of M and E peaks of *D. melanogaster* [31], most likely by promoting the siesta between them [32]. Therefore, a subgroup of the DN1_p_ has been named “siesta neurons”. We revealed interesting differences in PDP1 oscillations between the two strains and *D. melanogaster*.

In *D. melanogaster*, PDP1 oscillated in all clock neurons examined and under all photoperiods, albeit with reduced amplitude under photoperiods different than LD12:12, which is consistent with previous observations (figure 3*b*, tables 1, S4, S5) [30]. Under very long photoperiods (LD20:4) the PDP1 oscillations became bimodal and, most strikingly, very high in the DN_1_ (the siesta neurons) and LN_d_ (the E neurons) (figure 3*b*, tables 1, S4, S5). This matches the activity patterns under LD20:4, where the siesta was long and E activity most pronounced (figure 2*d*).

In the southern *D. littoralis* strain (Batumi), PDP1 cycling was found to be rhythmic in most clock neurons at LD8:16 and LD12:12, albeit the oscillations were not as synchronous as in *D. melanogaster* (figure 3*c*, tables 1, S4, S5). However, under the very long photoperiod (LD20:4), the LN_d_ and l-LN_v_ showed a bimodal pattern with high amplitude, while PDP1 levels became again high in the DN_1_ and lost rhythmicity in the s-LN_v_ (figure 3*c*; Table 1). Again, this staining pattern coincides with the locomotor activity patterns, which resembled those of *D. melanogaster*, but were generally noisier, particularly under the very long photoperiod (figure 2*d*).

**Table 1.** JTK-cycle analysis over 24h and 12h of PDP1 staining in the nuclei of different clock neurons.

| Strain | Photoperiod | Cluster | P value<br>(range = 24h) | P value<br>(range = 12h) |
| --- | --- | --- | --- | --- |
| *D. melanogaster* | LD8:16 | DN<sub>1</sub> | 1.61E-06 | 1 |
|  |  | LN<sub>d</sub> | 5.66E-07 | 0.63 |
|  |  | I-LN<sub>v</sub> | 2.25E-06 | 0.18 |
|  |  | s-LN<sub>v</sub> | 9.52E-06 | 0.24 |
|  | LD12:12 | DN<sub>1</sub> | 4.34E-12 | 0.19 |
|  |  | LN<sub>d</sub> | 9.60E-17 | 0.49 |
|  |  | I-LN<sub>v</sub> | 2.92E-12 | 0.20 |
|  |  | s-LN<sub>v</sub> | 3.70E-12 | 0.027 |
|  | LD20:4 | DN<sub>1</sub> | 0.40 | 0.00019 |
|  |  | LN<sub>d</sub> | 7.49E-06 | 0.0023 |
|  |  | I-LN<sub>v</sub> | 0.29 | 0.14 |
|  |  | s-LN<sub>v</sub> | 7.17E-06 | 0.01 |
| *D. littoralis*<br>Batumi | LD8:16 | DN<sub>1</sub> | 9.95E-05 | 0.01 |
|  |  | LN<sub>d</sub> | 0.10 | 0.35 |
|  |  | I-LN<sub>v</sub> | 0.42 | 0.52 |
|  |  | s-LN<sub>v</sub> | 4.44E-07 | 0.14 |
|  | LD12:12 | DN<sub>1</sub> | 0.008 | 1 |
|  |  | LN<sub>d</sub> | 0.002 | 0.77 |
|  |  | I-LN<sub>v</sub> | 0.003 | 1 |
|  |  | s-LN<sub>v</sub> | 4.10E-06 | 0.0020 |
|  | LD20:4 | DN<sub>1</sub> | 0.82 | 1 |
|  |  | LN<sub>d</sub> | 1 | 0.017 |
|  |  | I-LN<sub>v</sub> | 1 | 0.059 |
|  |  | s-LN<sub>v</sub> | 0.26 | 0.32 |
| *D. littoralis*<br>Kilpisjärvi | LD8:16 | DN<sub>1</sub> | 0.79 | 0.73 |
|  |  | LN<sub>d</sub> | 1 | 0.10 |
|  |  | I-LN<sub>v</sub> | 0.75 | 0.15 |
|  | LD12:12 | DN<sub>1</sub> | 0.30 | 1 |
|  |  | LN<sub>d</sub> | 0.10 | 0.03 |
|  |  | I-LN<sub>v</sub> | 1 | 0.99 |
|  | LD20:4 | DN<sub>1</sub> | 0.58 | 0.11 |
|  |  | LN<sub>d</sub> | 0.73 | 8.04E-06 |
|  |  | I-LN<sub>v</sub> | 0.81 | 1.69E-06 |

In the northern *D. littoralis* strain (Kilpisjärvi), we could neither see a clear staining of PDP1 nor PER in the s-LN_v_ (figure S3*g*). Moreover, under LD8:16, we detected no significant daily cycling of PDP1 in any of the clock neurons (Table 1), and if there were oscillations, these were largely out of phase with each other (figure 3*d*, table S4). Under LD12:12, the LN_d_ and l-LN_v_ oscillated in a daily manner, while the oscillations in the DN_1_ were not significant (Table 1). Under the very long photoperiod, the oscillations of the l-LN_v_, and LN_d_ became bimodal and of high amplitude (figure 3*d*). Indeed, JTK-cycle analysis revealed significant 12 h rhythms but no significant rhythms in the 24 h range (tables 1, Table **S**4, S5). In contrast to the southern *D. littoralis* strain, PDP1 levels in the DN_1_ were much lower and did not increase with increasing photoperiod in the northern strain (figure 3*d*, tables 1, S5). Again, this pattern of PDP1 cycling fits to the behavioural data, where the flies showed no clear organization in morning and evening activity under short days but exhibited morning and late evening activity with little siesta in between under longer days (figure 2*d*).

### 2.5. Latitude-dependent variations in PDF pattern and cycling

The most striking characteristic of the clock neuronal network in most of the *virilis* group strains studied so far is that the s-LN_v_ were not marked by antibodies against PDF [13,15–17,19]. Furthermore, most of these studies showed additional non-clock PDF-positive neurons in the dorsal brain of these species, which were recently shown to be CRZ-positive LNCs [18]. Here, we investigated whether these PDF staining patterns were conserved in the five *D. littoralis* strains stemming from different latitudes. We found that the s-LN_v_ terminals of the five strains were marked by anti-PDF to different degrees (figure 4*a*). While the more southern strains Batumi, Portugal and Zürich showed prominent staining in the s-LN_v_ terminals, this staining was very faint in the northern strain Raattama and completely absent in Kilpisjärvi as found previously. Another notable difference between the strains was the intensity of the PDF immunostaining in the arborizations of the CRZ neurons. While the Kilpisjärvi strain showed high PDF immunoreactivity in both somata and arborizations of the CRZ neurons, the PDF staining intensity in the arborizations of the CRZ neurons appeared to decrease with latitude (figure 4*a*).

**Figure 4.**
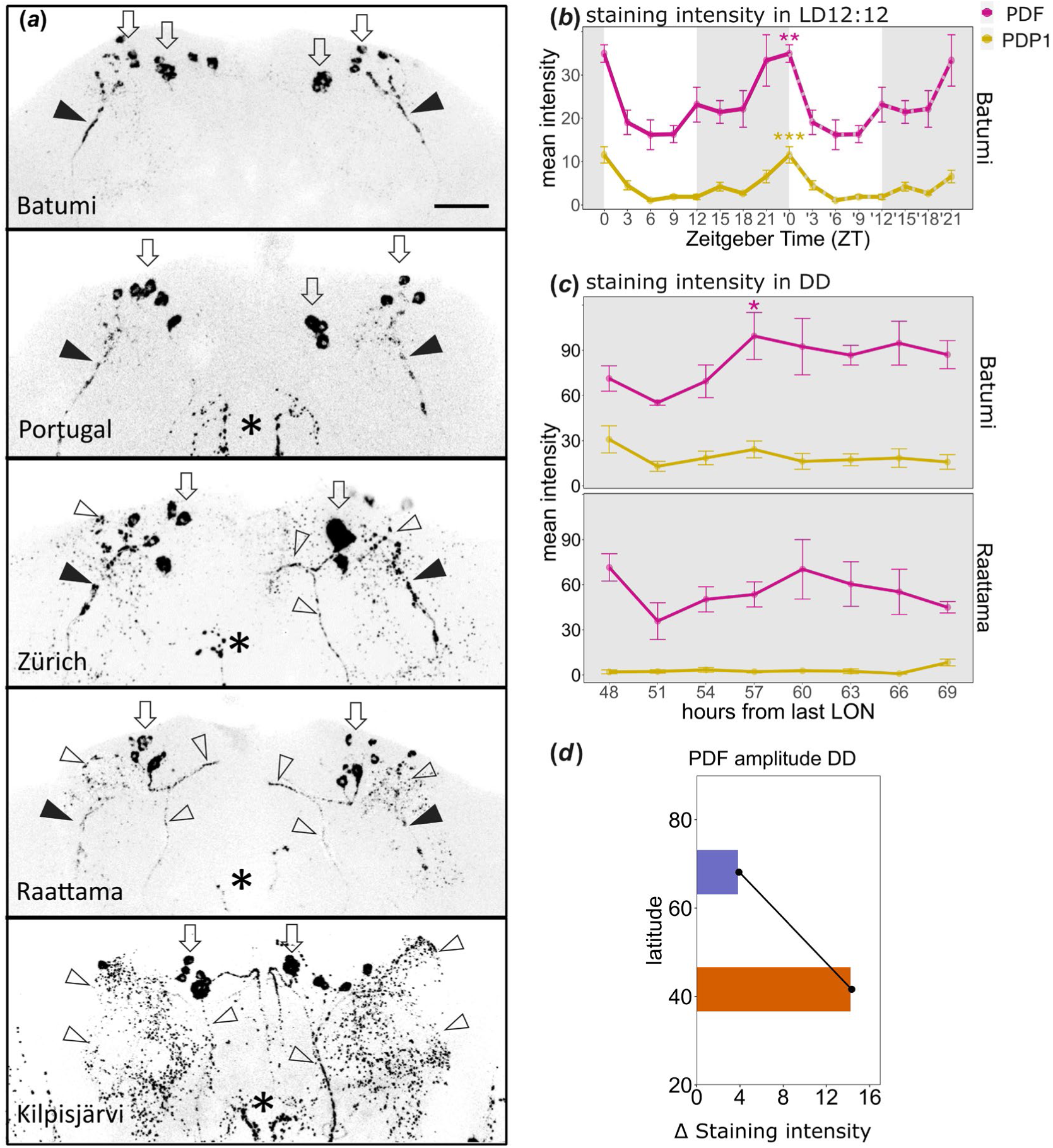
PDF staining in southern and northern *D. littoralis* strains and PDF/PDP1 oscillations in a northern and southern strain. (*a*) PDF staining in the brains of flies originating from different latitudes. The brains were stained within the first 2 hours after L_ON_ in LD20:4. Close arrowheads mark the terminals of the s-LN_v_, open arrowheads the arborizations of the CRZ neurons, the somata of which are indicated by open thick arrows. In the brain of the strains from Batumi, Portugal and Zürich, the PDF-positive s-LN_v_ arborizations were clearly visible, while they are only weakly stained in the northern strain Raattama and not visible at all in the strain from Kilpisjärvi. The somata of the CRZ neurons were always marked by anti-PDF, but their neurites appear more prominently stained in the northern strains. Asterisks mark fibers stemming from the l-LN_v_. Scale bar = 50µm. (*b*) Quantification of PDF and PDP1 immunostaining intensity in the brain of the southern *D. littoralis* strain from Batumi under LD12:12. The cycle of PDF (magenta) and PDP1 (yellow) were synchronized and peaked at ZT0 (Zeitgeber Time 0 = Lights on) (the data for PDP1 are the same as in figure 3c). (*c*) Upper panel, quantification of PDF and PDP1 immunostaining intensity in the brain of the southern *D. littoralis* strain from Batumi during the second day of DD after being entrained to LD12:12. The PDF staining intensity in Batumi was significantly cycling. Lower panel, quantification of PDF and PDP1 immunostaining intensity in the brain of the northern *D. littoralis* strain from Raattama during the second DD day. No significant cycling was found. *** p<0.001, ** p < 0.01, * p < 0.05, after JTK_CYCLE. (*d*) Amplitude of PDF cycling in the southern and northern strain plotted against latitude (table S6). The PDF cycling amplitude decreases with increasing latitude.

In *D. melanogaster*, PDF is essential for rhythmic behaviour under DD conditions [33], and it oscillates in a daily and circadian manner in the terminals of the PDF-positive s-LN_v_ with a maximum in the morning [34]. Therefore, we tested whether PDF also oscillates in the s-LN_v_ of the behaviourally rhythmic southern *D. littoralis* strain from Batumi. Under LD12:12, we found a significant cycling of PDF staining intensity in the s-LN_v_ terminals with a peak at L_ON_ (figure 4*b*, table S8). The PDF oscillation occurs in phase with the nuclear PDP1 oscillation that has already been shown in figure 3*c* and is again shown as a reference in figure 4*b*. This result indicates a rhythmic release of PDF from the s-LN_v_ terminals in the southern strain during LD cycles as it was shown in *D. melanogaster* [34].

To assess whether the cycling of PDF continues under DD conditions, we immunostained the brains of the flies of the behaviourally rhythmic *D. littoralis* strain Batumi and the behaviourally arrhythmic strain Raattama with antibodies against PDF and PDP1 every 3 hours on the second day of DD after being entrained to LD12:12 (figure 4*c*). We found that the molecular oscillations appear to continue in Batumi, although only PDF-cycling turned out to be significant by JTK_CYCLE analysis (Table S6-S8). In the northern strain Raattama, PDF did not significantly oscillate in the s-LN_v_ terminals (figure 4*c*; tables S6-S8), which fits well with its behavioural arrhythmicity under DD (figures 2; S1).

Remarkably, the amplitude of the PDF oscillation in the s-LN_v_ terminals follows the same general trend (figure 4*d*) as the behavioral phenotypes described above (figure 2), suggesting a possible causal effect.

### 2.6. Latitude-dependent seasonal changes in PDF and CRZ

Previous studies in *D. melanogaster* have shown that PDF is not only involved in the circadian control of behavioural rhythms, but most likely also in the seasonal control of reproductive dormancy [21,22]. The s-LN_v_ terminals of reproductive flies are significantly longer and more prominently marked by anti-PDF than those of dormant flies [21]. In high-latitude *D. littoralis* flies, these terminals are PDF-negative, but instead the neurosecretory CRZ neurons are stronger stained by the PDF antibody in reproductive flies (in LD20:4, 23°C) than in diapausing flies (in LD8:16, 10°C) (figure 5*b*) [18]. In contrast, CRZ levels are low in the same neurons in reproductive flies and high in diapausing flies (figure 5*b*) [18]. This suggests a putative antagonistic role of PDF and CRZ in the neurosecretory CRZ neurons in seasonal adaptation of northern *D. littoralis* flies.

**Figure 5.**
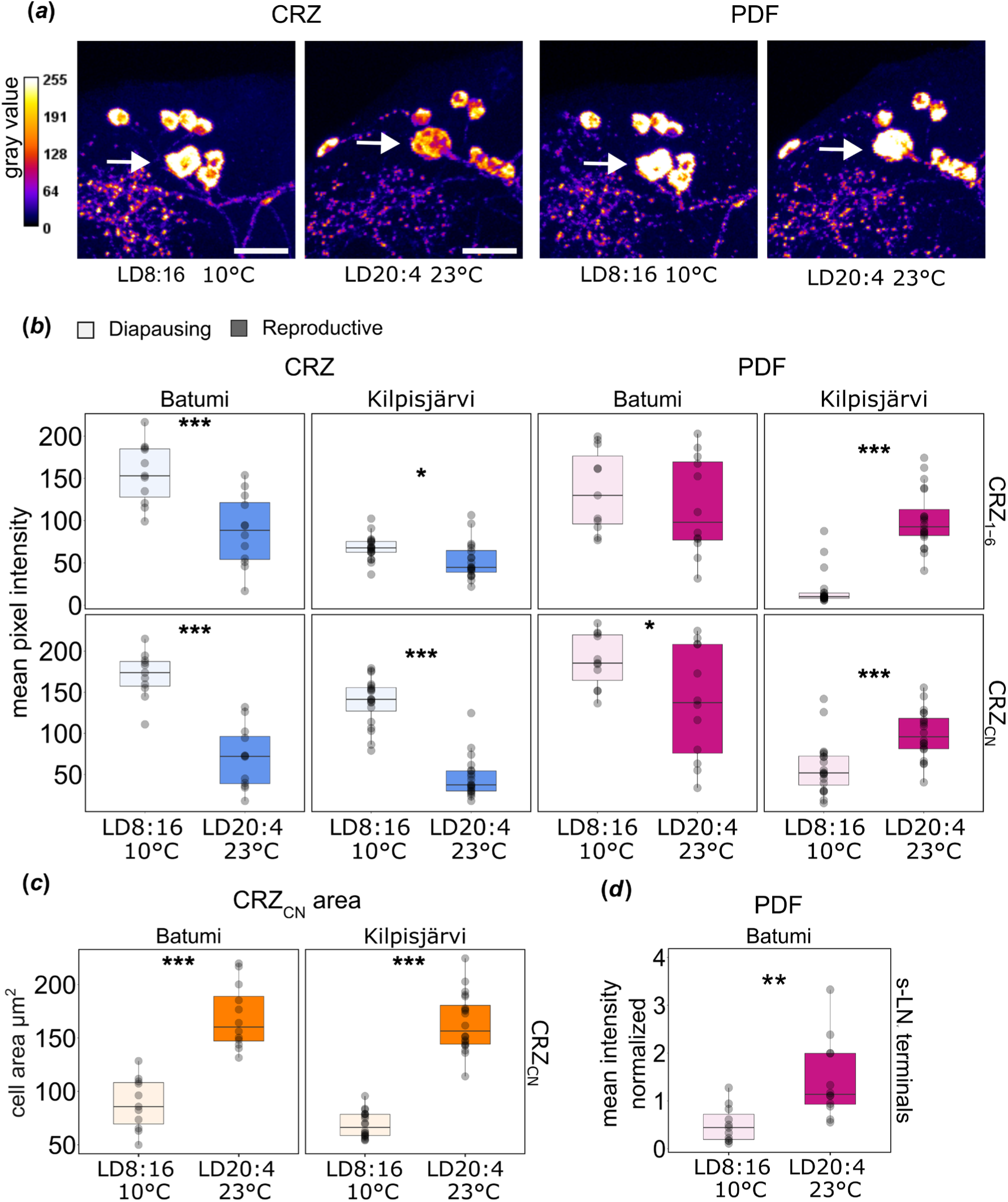
Quantification of CRZ and PDF staining intensity in the CRZ neurons of diapausing and reproductive *D. littoralis* from Batumi and Kilpisjärvi. (*a*) Grey value intensity of representative CRZ and PDF immunostaining in the CRZ neurons in the brain of the southern strain from Batumi. CRZ_CN_ are marked by a white arrow, they differ by size from the other CRZ_1-6_. (*b*) Quantification of the CRZ immunostaining intensity (left panel). In both CRZ clusters and both *D. littoralis* strains there was a significant decrease in the intensity under reproductive conditions (blue) compared to the diapausing conditions (light blue). The quantification of PDF immunostaining intensity (right panel) revealed no significant difference between the diapausing condition (light magenta) and the reproductive conditions (magenta) in the CRZ_1-6_ neurons of the southern strain of *D. littoralis* from Batumi. Nonetheless, a slight increase in PDF intensity was present in this species in the diapausing CRZ_CN_. In *D. littoralis* from Kilpisjärvi instead, the PDF staining intensity was significantly higher under reproductive conditions in both CRZ clusters. (*c*) The quantification of the cell body area of the CRZ neurons showed a significant increase in flies under reproductive conditions (orange) compared to diapausing conditions (light orange) in both strains tested. (*d*) The quantification of the PDF staining intensity in the s-LN_v_ terminals revealed an increase in the intensity when the flies from Batumi were exposed to reproductive conditions (magenta), compared to diapausing ones (light magenta). *** p<0.001, ** p<0.01, * p<0.05, after Wilcoxon test in CRZ intensity in CRZ_CN_ of Kilpisjärvi and in PDF intensity in Kilpisjärvi and t-test in the remaining ones. Scale bars 50 µm.

To judge the role of PDF in seasonal adaptation in a southern *D. littoralis* strain, we quantified the immunostaining intensity of PDF and CRZ in the CRZ neurons of the strain Batumi under diapause-inducing conditions (LD8:16, 10°C) and reproductive conditions (LD20:4, 23°C). These neurons can be subdivided by morphology into two groups, six smaller CRZ_1-6_ neurons and one bigger neurosecretory CRZ_CN_ neuron per hemisphere (figure 5*a* white arrow) [18,35], hence we measured separately the staining intensities of the two clusters. We compared directly the newly acquired data in Batumi to the previously published ones in Kilpisjärvi [18]. As previously found in the Kilpisjärvi strain, the CRZ immunostaining intensity in both CRZ clusters was also higher in the Batumi strain under diapausing conditions (figure 5*b*). However, the PDF intensity in the CRZ neurons of the Batumi strain was not significantly different between the two conditions in CRZ_1-6_ neurons and slightly increased in the CRZ_CN_ under diapausing condition (figure 5*b*). Next, we measured the size of the CRZ neurons somata and found that these were significantly larger under long summer days in both strains (figure 5*c*) suggesting that more peptides can be stored in the CRZ neurons somata when the flies are reproductive.

The present results indicate that CRZ plays a similar role in inducing diapause in both strains, while PDF in the CRZ neurons is playing a different role in the southern *D. littoralis* strain compared to the northern one. To test whether PDF from the s-LN_v_ terminals showed a seasonal variation in the southern *D. littoralis* strain, as it does in *D. melanogaster*, we quantified the PDF staining intensity in the PDF-positive s-LN_v_ terminals in Batumi under simulated winter and summer days. We found a significantly higher PDF staining in the s-LN_v_ terminals of reproductively active flies than in diapausing flies (figure 5*d*). This suggests that southern *D. littoralis* flies control diapause in a similar way as *D. melanogaster* flies control dormancy.

## 3. Discussion

A long-standing question in chronobiology is whether the circadian clock is involved in measuring day/night length to prepare organisms in good time for the coming winter and summer, as Bünning (1936) [7] hypothesized many years ago. Although many studies support such a role for the circadian clock [36–38], this is still under debate in insects for three main reasons: 1) Photoperiodic timing appears to have evolved differently from circadian timing in the pitcher plant mosquito, *Wyeomyia smithii* [39]. 2) Classical Nanda-Hamner experiments, which can prove the involvement of the circadian clock in measuring day or night length [40], give negative results for some insects including fruit flies [41,10,15]. 3) A knock-out of the clock gene *period* in *D. melanogaster* does not abolish the flies’ photoperiodic responses [42].

Pitcher plant mosquitoes have spread from south to north and from low to high elevations in eastern North America and therefore several populations stemming from different altitudes and latitudes can be investigated for diapause and clock characteristics [8,43]. In these populations, the critical photoperiod for the onset, maintenance and termination of diapause increased linearly with both altitude and latitude [44], but no such latitudinal clines have been observed in oscillation period of the circadian clock [39]. The authors speculate that this is because the photoperiodic timer and the circadian clock serve two very different functions with different requirements. While the photoperiodic timer must evolve rapidly in response to climatic changes, the circadian clock must be stable to maintain the coordination of daily rhythms. This is certainly true, but latitudinal clines in rhythm strength have also been observed in pitcher plant mosquitoes: as shown here for *D. littoralis*, the circadian clock of *W. smithii* weakens with increasing latitude [8].

More recently, experiments with *D. montana* even demonstrated latitudinal variations of Nanda-Hamner curves: the southern populations entered diapause only when the day lasted about 24h, while northern strains entered diapause even at Zeitgeber cycles much longer than 24h [41]. A possible explanation for this is that weak clocks can still stably entrain to very long Zeitgeber cycles and can therefore serve as time reference for measuring night length, while this is not the case for strong clocks which free-run under such conditions and therefore fail to report night length.

Finally, *D. melanogaster* flies stem from the tropics and do not show a strictly photoperiodic diapause, but rather a reproductive dormancy in response to stressful conditions such as cold temperatures [45]. This makes them not suitable to prove the involvement of the clock gene *period* in photoperiodic time measurements.

### 3.1. Hypothetic link between the robustness of the clock, phases of M_peak_ and E_peak_, and critical night length for diapause induction

Flies originating in northern latitudes face very rapid changes in photoperiod that can best be tracked by weak plastic clocks. Such plastic clocks can not only better follow very long Zeitgeber cycles (see above), but also very long photoperiods [11,19,46]. Indeed, as latitude increased, we found a stronger coupling of M_peak_ and E_peak_, to the beginning and end of the photoperiod. We hypothesize that the circadian clock of the common ancestor of the *virilis* group was like that of the southern strains of *D. littoralis* and that, being weaker than the clock of *D. melanogaster*, it allowed this group to invade more northern latitudes. Their clocks, once they reached northern latitudes, may have weakened further, becoming fast-dampening clocks. In parallel, the tracking of dawn and dusk by the weak M and E oscillators improved further.

The next question is whether the different phases of M_peak_ and E_peak_ affect the CNL for diapause induction in southern and northern *D. littoralis* strains. Assuming that the time interval (phase relationship) between E_peak_ and M_peak_ codes for night length as predicted by the Internal Coincidence Model [47,48], this time interval is very short for northern flies under LD20:4 and longer for southern flies, as the latter cannot couple their E_peak_ and M_peak_ to dusk and dawn under extreme photoperiods (see figure 2*e*). It is not difficult to imagine that, in the northern strains, the short night signal also sets the CNL to a short night, while the longer night signal of the southern flies leads to a longer CNL. Alternatively, the CNL could also be influenced by the time interval between the beginning of the night and the M_peak_, which is also very short for the northern flies under LD 20:4 and significantly longer for the southern flies. This measurement of night length would correspond to the quantitative version of the External Coincidence Model, which generally fits flies better than the internal coincidence model [2,11,36,41]. In this model, the assessment of night length is based on the accumulation of a "diapause" substance during darkness. As soon as this substance reaches a certain reference threshold, diapause is initiated. In the northern strains the critical threshold might be reached earlier because diapause-inhibiting factors such as PDF are reduced (see below).

### 3.2. Molecular and neuronal basis for the latitudinal adaptation of the circadian clock

At the molecular level, we found differences in the daily oscillations of several clock components in southern and northern *D. littoralis* strains that are consistent with the observed behaviour at different photoperiods and constant darkness. In the southern species, the daily PDP1 oscillations were robust and highly synchronous between the assessed clock neurons under short days while they were barely significant and quite out of phase in the northern *D. littoralis* strain. However, during long days, significant 12-h rhythms occurred in a few clock neurons in the northern *D. littoralis* strain, which might drive the observed bimodal activity pattern with M_peak_ and E_peaks_. Furthermore, PDP1, PER and PDF were either completely absent or not oscillating in the s-LN_v_ of the northern strain. This could explain the flies’ behavioural arrhythmicity under DD conditions since these neurons are important for robust activity rhythms under DD conditions in *D. melanogaster* [33,49,50].

### 3.3. The neuropeptide PDF as a putative link between the circadian and seasonal clock

PDF is not only involved in the circadian clock of several insects [51–53] and in behavioural plasticity [54], but it has an additional role in photoperiodism. Blow flies (*Protophormia terraenovae*) cannot discriminate between long and short days when their PDF-positive clock neurons are ablated [55]. *Culex pipiens* mosquitoes, need PDF to remain reproductive under long day conditions [56], while Linden bugs (*Pyrrhocoris apterus*) [51] and brown-winged green bugs (*Plautia stali*) [57] need PDF to enter diapause under short days. A role of PDF in photoperiodism was most recently also suggested for the pea aphid *Acyrtosiphon pisum* [58,59]. In *D. melanogaster*, PDF released from the s-LN_v_ acts on the insulin-producing MNCs and inhibits diapause [21,22]. Thus, PDF is needed to keep the flies in the reproductive state similar to mosquitoes. The same seems to be true in the southern *D. littoralis* strain Batumi, which shows enhanced PDF staining in the s-LN_v_ terminals under long photoperiods as previously shown in *D. melanogaster* [21]. Thus, we propose that the diapause-inhibiting effect of PDF is conserved in *D. littoralis*, although the way in which PDF acts differs in the southern and northern strains. The northern strain from Kilpisjärvi lacks PDF in the s-LN_v_ terminals and consequently also the diapause-inhibiting effect via the MNCs. This might facilitate the induction of diapause in the northern flies already at longer photoperiods and result in shorter CNLs (see above). On the other hand, under reproductive conditions, the northern strains strongly express PDF in the CRZ neurons and in their fibres running to the *corpora cardiaca*. There, PDF may act on PDF receptors on adipokinetic hormone-expressing cells and increase metabolic activity [60]. PDF may even be released via the *corpora cardiaca* into the circulation and could target peripheral tissues that express the PDF receptor [61,62]. In northern *D. littoralis*, *Pdf* transcript and PDF peptide levels increase in the CRZ-positive neurons under summer conditions [18]. This could mean that PDF stemming from the CRZ-positive neurons may have overtaken the role of clock neuron-derived PDF in promoting the flies’ reproductivity in summer, although this way seems clearly less effective than the way via the insulin-producing MNCs. CRZ on the other hand, is a good candidate for inducing diapause when days shorten in autumn, and this appears to be similarly true in southern and northern strains as we show here. In summary, northern and southern *D. littoralis* appear to differ in the way by which PDF keeps them in the reproductive state, but not in the way CRZ promotes diapause. Nonetheless, PDF and CRZ are clearly not the only neurohormones involved in diapause regulation. More studies are needed to unravel the influence of the endogenous clocks on the neurohormonal system in *D. littoralis* and other insects.

### 3.4. Conclusions

Our study extends the results of previous studies that found a latitudinal cline in eclosion rhythms of *D. littoralis* to the adult locomotor activity. We found that several behavioural, neuronal and molecular features of the adult circadian clock show latitudinal clines that are consistent with the latitudinal cline of the photoperiodic diapause, suggesting the potential involvement of the circadian timekeeping system in seasonal adaptation. In particular, differences in the expression pattern and oscillation of the neuropeptide PDF in the fly brain could be responsible for the observed differences in seasonal diapause timing between *D. littoralis* populations living at high- and low-latitudes. This is an important step in revealing the role of the circadian clock in the seasonal response of insects.

## 4. Material and methods

### 4.1. Fly strains and husbandry

The different *D. littoralis* strains were collected over different years throughout Europe. The strains collected in Kilpisjärvi (69°N) and Rovaniemi (66°N) (Finland), Zürich (47°N) and Ticino (46°N) (Switzerland), in Batumi (41°N) and Khobi (42°N) (Georgia), were first collected and described in the study by Lankinen [12]. The Portugal strain was collected in Portugal (41°N) in the year 2000. The strains collected in Raattama (68°N), and Petäjävesi (62°N) (Finland), were kindly provided by Maaria Kankare and were collected in 2020 and 2021, respectively (Figure 1*a*). These strains were reared at 23°C under Light-Dark cycles (LD) 20:4 with 60-70% RH. The *D. melanogaster* stock used was the Canton-S wild-type line [63] and was reared under LD12:12 at 25°C with 60% RH. All flies were fed on standard cornmeal/agar medium with yeast.

### 4.2. Estimating the critical night length

These experiments were performed in Oulu, while all the other experiments presented in this study were performed in Würzburg. The flies were maintained at 16°C, LL and fed on malt medium [64]. Different groups of freshly eclosed female flies were entrained to different photoperiods at 16°C. Their ovaries were dissected after three weeks of entrainment; following Lankinen et al. (2021) [41] the flies were scored as reproductive in the presence of fully developed eggs and diapausing when little to no yolk was present. Photoperiodic response curves (PPRCs) of the different strains were calculated using the function *drm* (dose response models) from the R package drc (dose response curve, v4.2.3; [41,65]). The Critical night length (CNL) is found at the point of inflection of the PPRC (ED50 = 50% of the flies are diapausing) (see figure 1*c*).

### 4.3. Locomotor activity recording

The locomotor activity of male flies aged between 3 to 7 days post eclosion, was recorded with the *Drosophila* Activity Monitor (DAM) system (Trikinetics, Princeton, MA, USA). Single flies were placed in glass tubes with a diameter of 0.5 cm for *D. melanogaster* and 0.7 cm for *D. littoralis*. The tube was filled on one end with food consisting of 4% saccharose and 2% agar in water, while the other end was blocked by a foam plug. The activity was recorded for approximately 10 days under different LD cycles: LD8:16, LD12:12 or LD20:4 at 20°C. Afterwards, the flies were exposed to constant darkness (DD) to assess their rhythmicity under constant conditions for at least 9 days. Of the 10 days recorded in LD, only the last 5 were used for the calculation of average activity profiles and activity peaks, while circadian rhythmicity, period length and power in DD were calculated for the first 9 days of constant darkness.

### 4.4. Neuropeptide expression in diapausing and reproductive flies

To quantify the staining intensity of the neuropeptides PDF and CRZ in the CRZ neurons of reproductive and diapausing flies, 60 virgin females of the Batumi strain were collected and exposed to diapause-inducing conditions (LD8:16, 10°C) or reproductive conditions (LD20:4, 23°C). After three weeks, the flies were collected at ZT1 (Zeitgeber Time1, which corresponds to one hour after lights-on) and fixed as described below. The flies’ ovaries were dissected in a phosphate-buffered saline solution (PBS) and their reproductive state was determined as described above. Afterwards, the brains were dissected and further stained as described below.

### 4.5. Fluorescent immunohistochemistry and microscopy

Immunohistochemistry was performed on male flies between 3 to 7 days post eclosion following the protocol from Schubert et al. (2018) [66]. We immunostained the flies’ brains to define the anatomical position and staining intensity of the circadian clock proteins PERIOD (PER) and Par Domain Protein 1 (PDP1) and the circadian clock neuropeptide Pigment Dispersing Factor (PDF) in different strains of *D. littoralis*. Flies were fixed for 3.5 h in 4% paraformaldehyde dissolved in PBS with 0.5% Triton X100 (PBST) added. The flies were then dissected after 3 washes of 10 min with PBS. Subsequently the brains were blocked in a 5% Normal goat serum (NGS) solution in PBST 0.5% (this was substituted by Normal donkey serum, NDS, when the primary antibody solution contained anti-PER-goat) overnight at 4°C. The brains were then incubated in the primary antibody mix (see Table 2 for information on the antibodies) which consisted of the primary antibodies with 5% NGS or NDS and 0.02% NaN_3_ in PBST 0.5% for 2 days at 4°C and one day at Room Temperature. After 6 PBST washes of 10 min, the brains were incubated overnight at 4°C in the secondary antibody mix (see Table 2) consisting of the secondary antibodies with 5% NGS (or NDS) in PBST 0.5%. Lastly, the brains were embedded with Vectashield (Vector Laboratories, Burlingame, CA, USA) on microscope slides with ∼150 µm spacer. The images were acquired with a Leica SPE confocal microscope (Leica Microsystems, Wetzlar, Germany), equipped with a photomultiplier tube and solid-state lasers (488, 532, and 635 nm) for excitation. Pictures were scanned with a 20-fold magnification glycerol immersion objective (HC PL APO; Leica Microsystems). The confocal stacks had a 2 μm z-step size and either 1024 × 1024 pixels for the quantifications (figures 4,5) or 2048 x 2048 (figure S3) with a pixel size of 537 × 537 nm and a voxel size of 0.537 × 0.537 × 2 μm. The images were further processed using Fiji (v 1.53t, [67]). Brightness and contrast were eventually changed linearly.

**Table 2.**
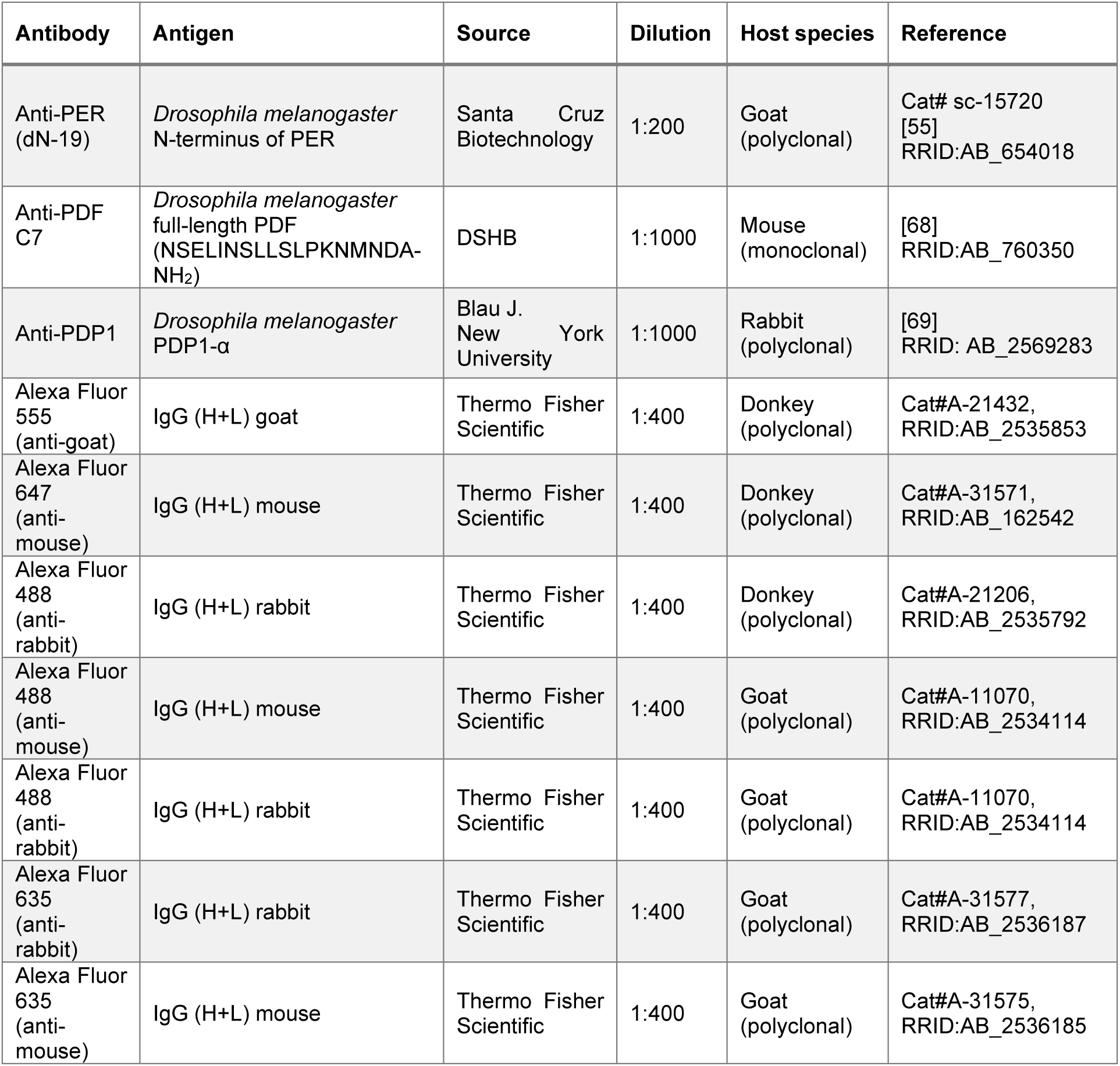
Primary and secondary antibodies used.

### 4.6. Quantification of immunostaining intensity

All the samples quantified were stained simultaneously under the same conditions, pictures were then acquired with the same settings. The quantification was carried out in Fiji on raw pictures. Since different types of signals were quantified, we used different methods for each quantification. To quantify the nuclear PDP1 staining intensity, a 3×3 px Region of Interest (ROI) was generated with the Fiji Rectangle selection tool on the brightest focal plane of the nucleus. To quantify the PDF staining intensity in the s-LN_v_ terminals, the ROIs were set on the 10 pixels with the highest intensity. To quantify the PDF and CRZ staining intensity in the CRZ neurons in the diapausing conditions, the ROIs were generated around the whole cell body. For all the different quantifications, the mean grey value of the ROI was then subtracted by the background intensity ROI value which was set near the quantified region and with a comparable ROI area. Finally, for quantifying the PDF staining intensity in the PDF terminal of Batumi flies kept under long and short days, we used the method described in Hermann-Luibl et al. (2014) [70]. First, 8 to 10 confocal stacks containing the s-LN_v_ terminals were overlayed as a maximal projection, using the function *Stacks Z Project…* and *Max Intensity* as projection type in Fiji. Second, the PDF-staining in the CRZ-neurons had to be removed. For this purpose, the brains had been fluorescently immunostained by anti-CRZ and anti-PDF, and the CRZ-channel was subtracted twice from the PDF-channel in ImageJ (*Process › Math › Subtract…).* This reliably eliminated the PDF-staining in the CRZ arborizations, but in rare cases some staining in the cell bodies remained, which was removed manually. In some cases, some PDF-staining in the posterior optic commissure or the optic lobes was visible, which was also removed manually. In the next step, the background was subtracted, so that everything in the picture remained black, except the PDF-terminals. Subsequently, the pixel intensity of the entire picture was measured with the tool *Analyze > Measure* in Fiji.

### 4.7. Statistical analysis and figures

Except for the actograms (Figure S1), which were generated with ActogramJ (a Fiji plugin) [71], all other graphs were generated with R (v4.2.3) via Rstudio (v2023.09.1+494), using the ggplot2 package [72]. The locomotor activity and sleep graphs were generated using the Rethomics framework [73]. The sleep amount was calculated with the function *sleep_dam_annotation* which scores as asleep individuals which are immobile for a minimum of 5 minutes. The average locomotor activity was calculated starting from the averaged locomotor activity of single flies over the last 5 days of LD, this was subsequently averaged among all flies and smoothed with a moving average filter over 11 of the 1 min values. The latitude map plot was generated with the packages rnaturalearth [74] and sf [75]. Further modifications to the figures were made using Inkscape (v1.3).

The photoperiod lengths recordings (figure 1*b*) were collected from: Open meteo > historical weather over 4 years, from 2019 to 2023 (https://open-meteo.com/en/docs/historical-weather-api).

The statistical analysis was performed in R. In general, all data were tested for normal distribution using Shapiro’s normality test and for homogeneous variance using Levene test. Based on the results of these tests, different approaches were taken, as described in the images caption. To compare the sleep during the light phase, the morning and evening peaks between strains, a Kruskal-Wallis test was performed followed by pairwise comparison using the Wilcoxon rank sum test. The Lomb-Scargle periodograms were generated using the rethomics framework [73]. The cycle analyses were performed by JTK_CYCLE [76] using the R package DiscoRhythm (v1.14.0) [77]. The single ROI intensities were averaged per brain and the analysis was run on the averaged brain values.

The evening peak quantification was carried out as described in Vaze et al., 2023 [11]. Briefly, the activity of single flies was averaged through 5 LD days, and the data was duplicated to obtain a double plot. The activity was then smoothed by local regression with a 0.02 span and finding the max values of the data with a rolling filter over 11 data points. The peaks were then found at the intersection points between the curve generated and the original activity data.

## Data accessibility

Raw data and data analysis are available in the supplementary section.

## Authors’ contributions

G.M., P.L., and E.B. performed and analysed the experiments, C.H.-F., and P.L. designed the study, G.M., E.B. and C.H.-F. wrote the manuscript with help from P.L.

## Acknowledgements

We would like to thank Pamela Menegazzi and Ina Götzelmann for their significant help in the quantification of PDP1 staining in *D. melanogaster*. We would also like to thank Jan Veenstra and Justin Blau for donating the anti-CRZ and anti-PDP1 antibodies, Maaria Kankare for donating the northern *D. littoralis* strains from Raattama and Petäjävesi, and Chedly Kastally for his help with the code for the CDL analysis. In addition, we are thankful to Sabrina Ender and Tom Viguir for their help in the locomotor activity recording and in staining of different *D. littoralis* strains brains.

## Supplementary information

**Figure S1.**
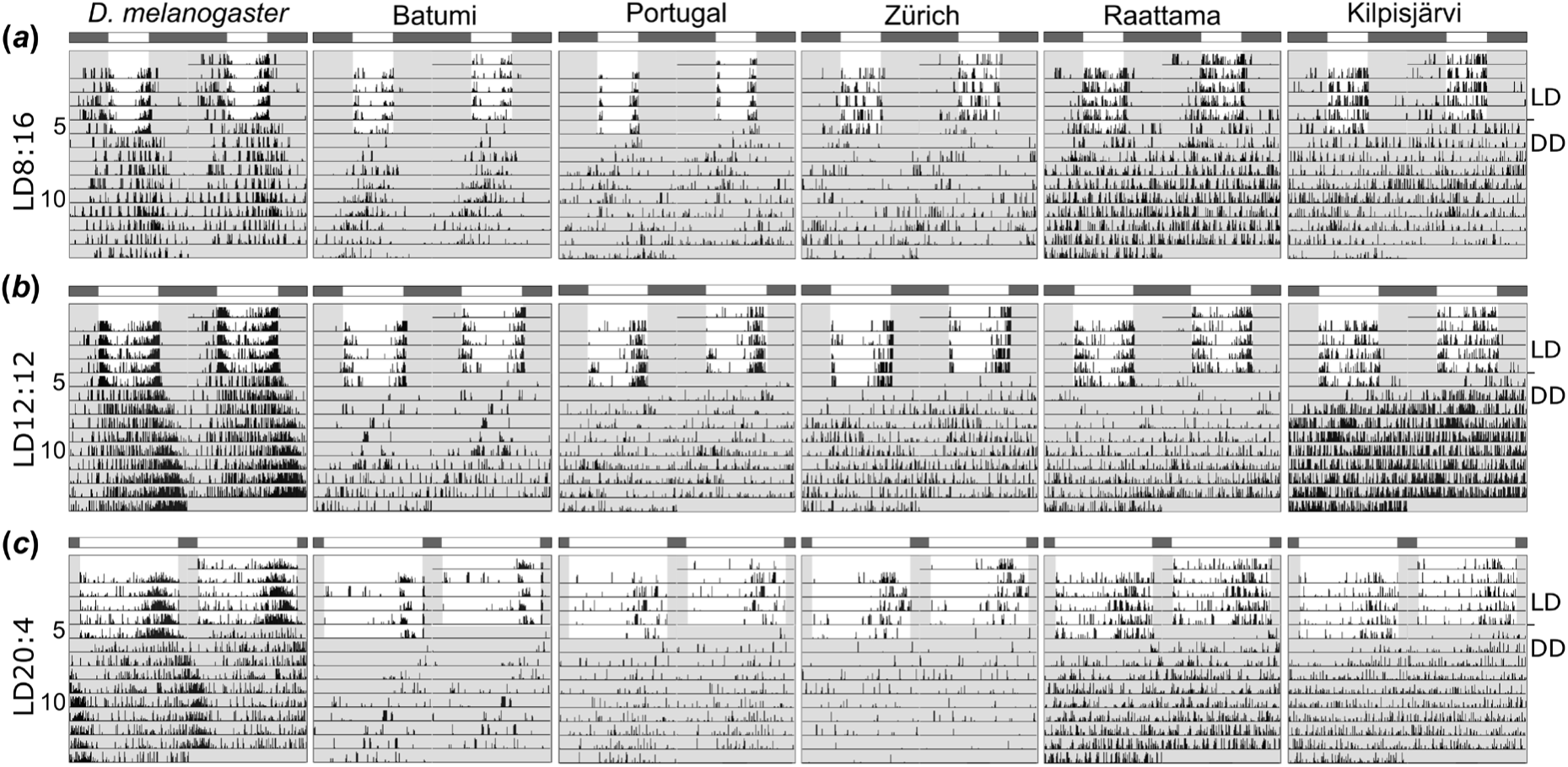
Double plots of representative actograms of individual *D. melanogaster* flies and *D. littoralis* flies from different latitudes under photoperiods of LD8:16, LD12:12, and LD20:4. Only the last five days of LD and the first 9 days of DD are shown. The light bars above the plots represent the photoperiod at which the flies were entrained for the first part of the experiment, from the sixth plotted day the flies were released in DD (the dark phases are shaded in grey). (*a*) Under LD8:16, the flies from the south including *D. melanogaster* showed a clear bimodal activity and remained rhythmic in DD, while the *D. littoralis* fly from Zürich and the two northern flies from Raattama and Kilpisjärvi lacked bimodality. Once released in DD the fly from Zürich remained weakly rhythmic while the northern flies became arrhythmic. (*b*) Under LD12:12, all flies except the fly from Kilpisjärvi showed bimodal activity. In DD, the *D. melanogaster* fly was strongly rhythmic, while the *D. littoralis* flies from Batumi, Portugal and Zürich remained weakly rhythmic and the northern flies became again arrhythmic. (*c*) Under LD20:4, the *D. melanogaster* fly showed weak delayed morning activity and robust advanced evening activity, while the southern *D. littoralis* flies and the fly from Zürich showed only evening activity. The two northern *D. littoralis* flies exhibited weak early morning activity and prominent late evening activity. In DD, the *D. melanogaster* fly showed robust rhythmicity, while the southern *D. littoralis* flies were weakly rhythmic and the flies from Zürich, Raattama and Kilpisjärvi were arrhythmic. LD = light-darkness cycles. DD = constant darkness.

**Figure S2.**
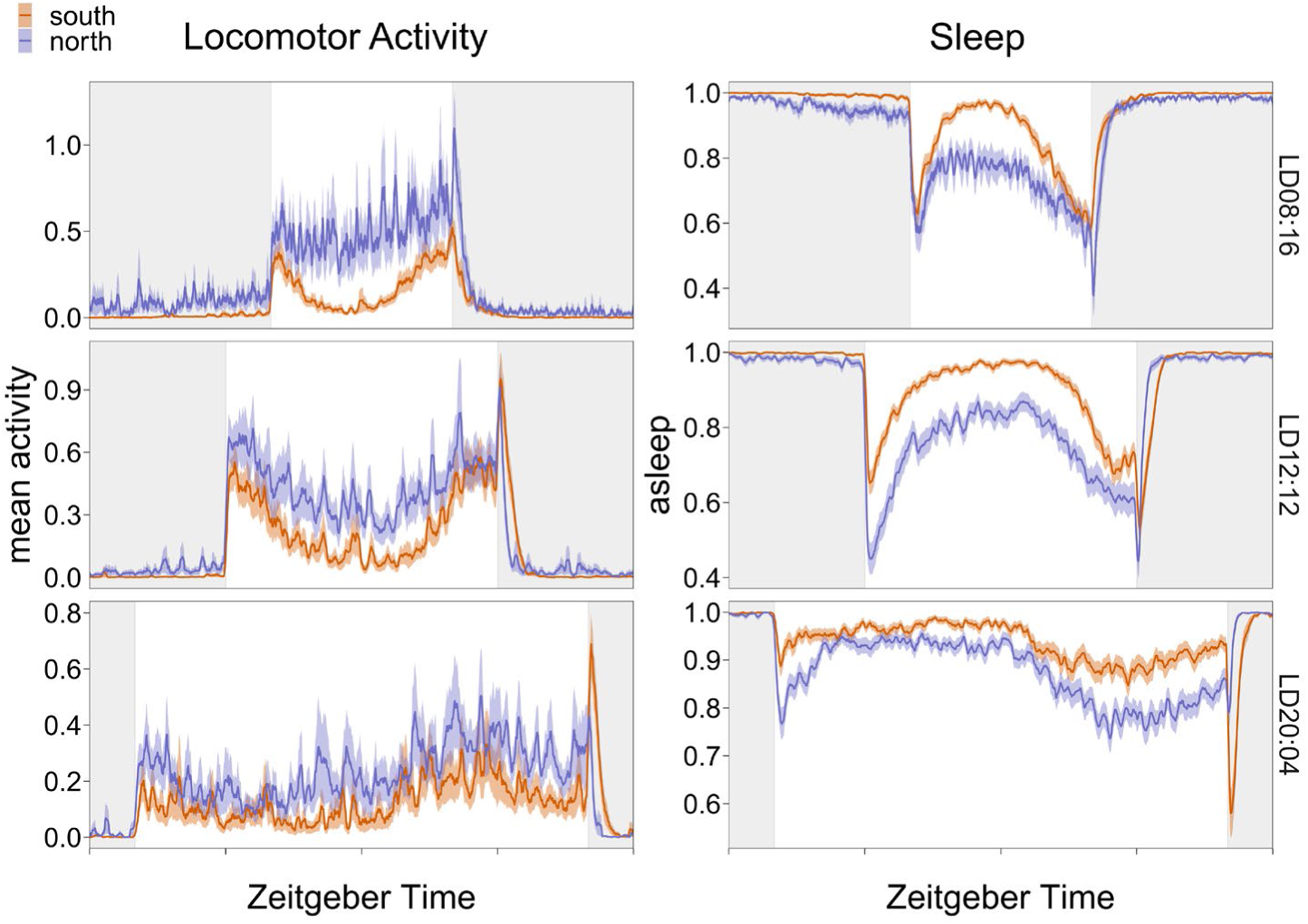
Mean activity and sleep averaged through northern strains (blue) and southern strains (orange).

**Figure S3.**
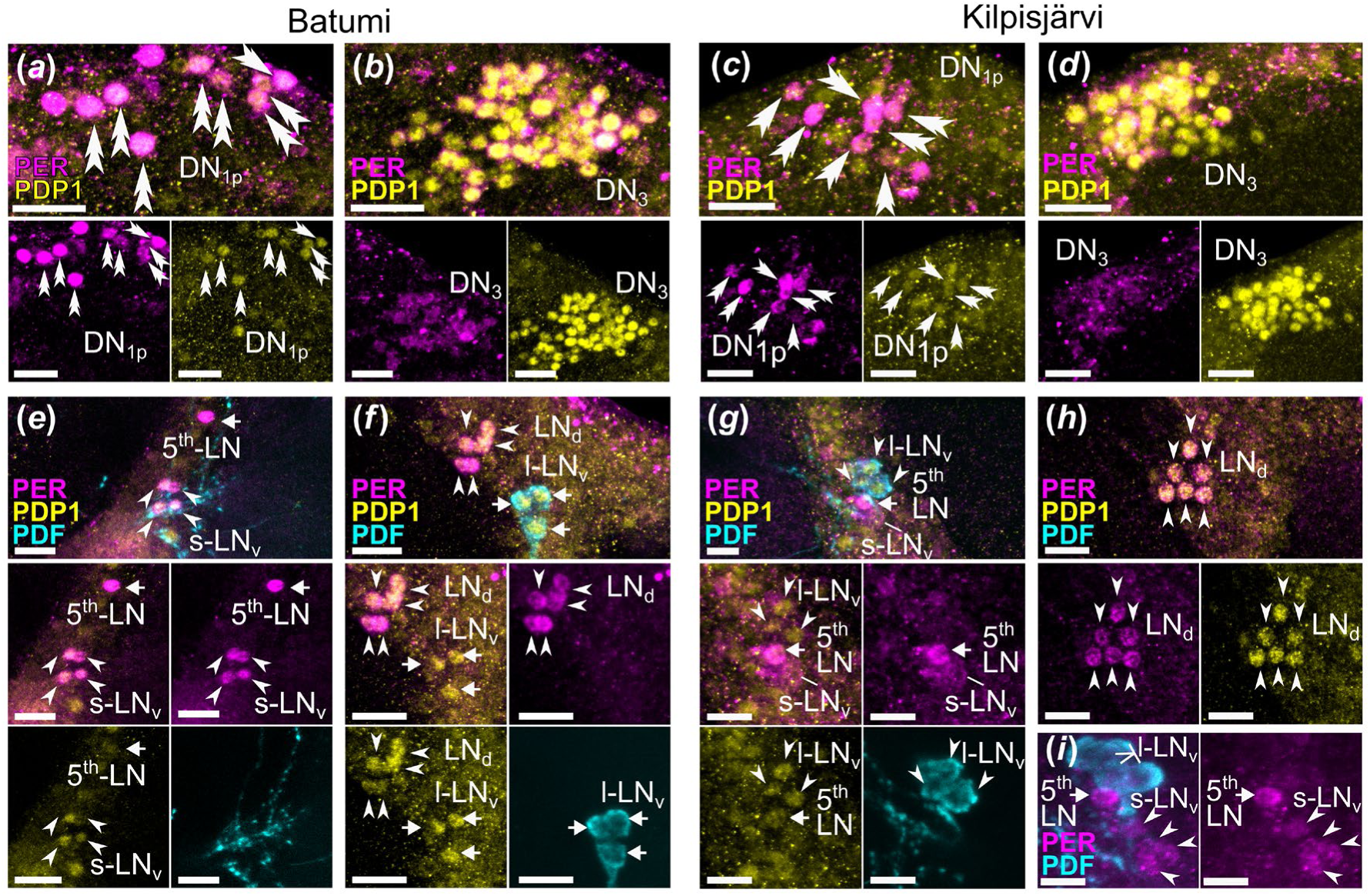
PDP1 and PER double-labelling in the brain of one southern and one northern strain of *D. littoralis* at ZT0. (*a*) PDP1 and PER colocalized in some DN_1p_ in the brain of the southern strain from Batumi (double arrow). (*b*) PDP1 and PER colocalized in most DN_3_ in Batumi. (*c*) PDP1 and PER colocalized in some DN_1p_ in the brain of the northern strain from Kilpisjärvi (double arrow). (*d*) PDP1 and PER colocalized in some DN_3_ in Kilpisjärvi. (*e*) PER and PDP1 colocalized in the 5^th^-LN_v_ (arrow) as well as in the s-LN_v_ (arrowheads) in Batumi. (*f*) PER and PDP1 colocalized in the LN_d_ (arrowheads), but not in the l-LN_v_ which showed only the PDP1 immunostaining (arrows) in Batumi. (*g*) PER and PDP1 were colocalizing in the 5^th^-LN (arrow), only PDP1 was detected in the l-LN_v_ (arrowheads), while in the s-LN_v_ PER was present at an extremely low level (line). (*h*) PER and PDP1 were colocalizing in the LN_d_ (arrowheads) of the Kilpisjärvi strain. (*i*) PER was not detected in the l-LN_v_ but was present in the 5^th^-LN (arrow) and faintly in the s-LN_v_ in the Kilpisjärvi strain. Scale bar = 50µm in A-D, J-L; 20µm in E-I.

**Table S1.** CNL and CDL in different strains of *D. littoralis* originating from different latitudes.

| CNL ±<br>Standard error | CDL | strain | latitude |
| --- | --- | --- | --- |
| 4.08 ± 0.67 | 19.92 | Kilpisjärvi | 69 |
| 4.80 ± 1.20 | 19.20 | Raattama | 68 |
| 5.64 ± 0.94 | 18.36 | Rovaniemi | 66 |
| 6.82 ± 1.33 | 17.18 | Petäjävesi | 62 |
| 7.50 ± 0.53 | 16.47 | Zürich | 47 |
| 11.41 ± 1.93 | 12.59 | Ticino | 46 |
| 10.23 ± 0.76 | 13.77 | Khobi | 42 |
| 10.35 ± 1.12 | 13.65 | Portugal | 41 |
| 11.48 ± 0.90 | 12.52 | Batumi | 41 |

**Table S2.** Number of flies tested in LD conditions.

| Strain | Photoperiod | N flies |
| --- | --- | --- |
| Canton-S | LD08:16 | 32 |
|  | LD12:12 | 46 |
|  | LD20:04 | 31 |
| Batumi | LD08:16 | 31 |
|  | LD12:12 | 56 |
|  | LD20:04 | 32 |
| Portugal | LD08:16 | 28 |
|  | LD12:12 | 30 |
|  | LD20:04 | 28 |
| Zürich | LD08:16 | 26 |
|  | LD12:12 | 33 |
|  | LD20:4 | 35 |
| Raattama | LD8:16 | 43 |
|  | LD12:12 | 33 |
|  | LD20:4 | 50 |
| Kilpisjärvi | LD8:16 | 31 |
|  | LD12:12 | 32 |
|  | LD20:4 | 27 |

**Table S3.**
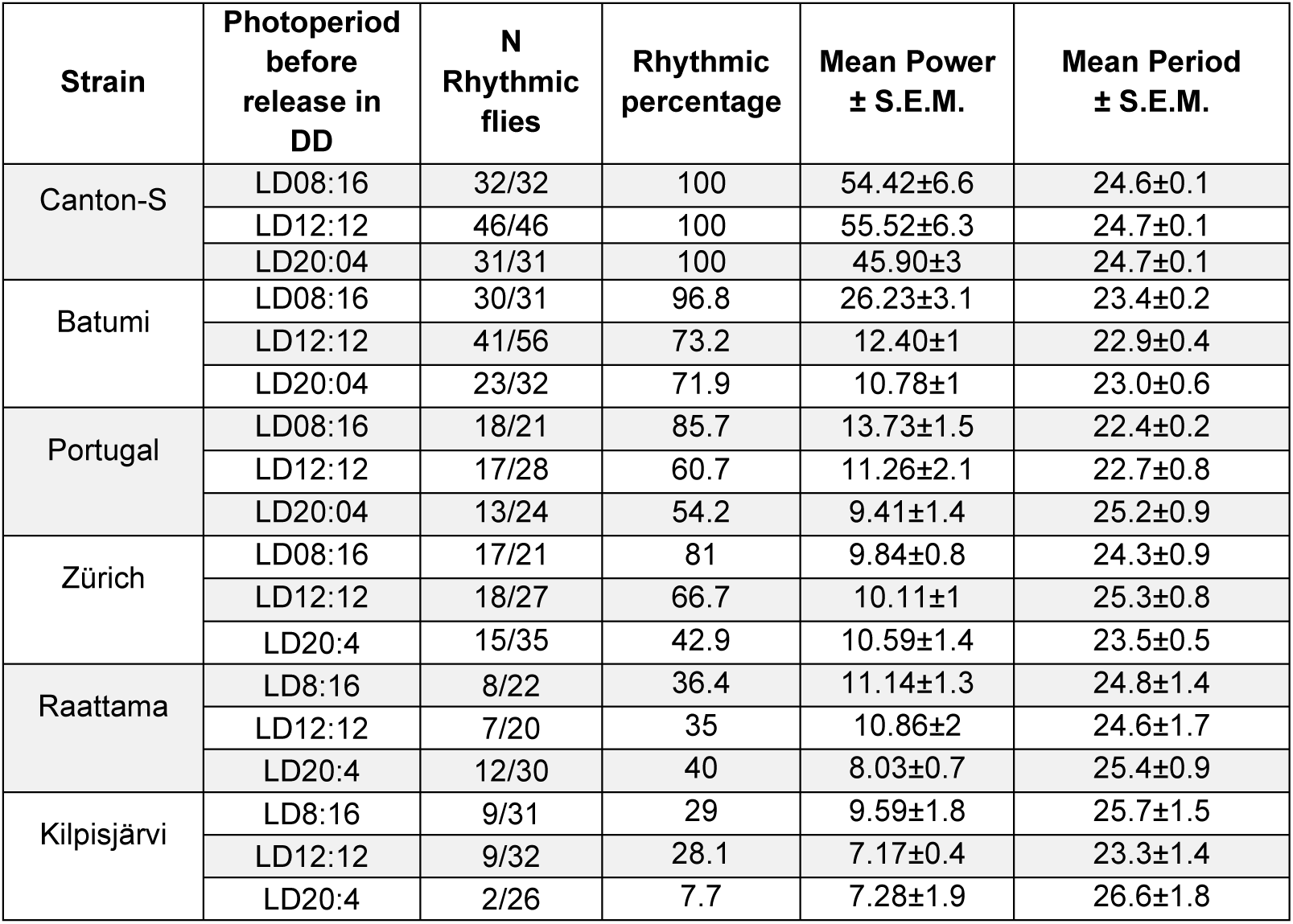
Lomb-Scargle periodogram in DD: population rhythmicity, power and period with p<0.05.

| Strain | Photoperiod<br>before<br>release in<br>DD | N<br>Rhythmic<br>flies | Rhythmic<br>percentage | Mean Power<br>± S.E.M. | Mean Period<br>± S.E.M. |
| --- | --- | --- | --- | --- | --- |
| Canton-S | LD08:16 | 32/32 | 100 | 54.42±6.6 | 24.6±0.1 |
|  | LD12:12 | 46/46 | 100 | 55.52±6.3 | 24.7±0.1 |
|  | LD20:04 | 31/31 | 100 | 45.90±3 | 24.7±0.1 |
| Batumi | LD08:16 | 30/31 | 96.8 | 26.23±3.1 | 23.4±0.2 |
|  | LD12:12 | 41/56 | 73.2 | 12.40±1 | 22.9±0.4 |
|  | LD20:04 | 23/32 | 71.9 | 10.78±1 | 23.0±0.6 |
| Portugal | LD08:16 | 18/21 | 85.7 | 13.73±1.5 | 22.4±0.2 |
|  | LD12:12 | 17/28 | 60.7 | 11.26±2.1 | 22.7±0.8 |
|  | LD20:04 | 13/24 | 54.2 | 9.41±1.4 | 25.2±0.9 |
| Zürich | LD08:16 | 17/21 | 81 | 9.84±0.8 | 24.3±0.9 |
|  | LD12:12 | 18/27 | 66.7 | 10.11±1 | 25.3±0.8 |
|  | LD20:4 | 15/35 | 42.9 | 10.59±1.4 | 23.5±0.5 |
| Raattama | LD8:16 | 8/22 | 36.4 | 11.14±1.3 | 24.8±1.4 |
|  | LD12:12 | 7/20 | 35 | 10.86±2 | 24.6±1.7 |
|  | LD20:4 | 12/30 | 40 | 8.03±0.7 | 25.4±0.9 |
| Kilpisjärvi | LD8:16 | 9/31 | 29 | 9.59±1.8 | 25.7±1.5 |
|  | LD12:12 | 9/32 | 28.1 | 7.17±0.4 | 23.3±1.4 |
|  | LD20:4 | 2/26 | 7.7 | 7.28±1.9 | 26.6±1.8 |

**Table S4.** PDP1 cycling, JTK_CYCLE analysis over a 24h period.

| Strain | Photoperiod | Cluster | Acrophase | Amplitude | P value | Q value | Mean |
| --- | --- | --- | --- | --- | --- | --- | --- |
| *D. melanogaster* | LD8:16 | DN<sub>1</sub> | 16.5 | 14.95 | 1.61E-06 | 1.61E-06 | 16.41 |
|  |  | LN<sub>d</sub> | 19.5 | 14.98 | 5.66E-07 | 5.66E-07 | 19.9 |
|  |  | I-LN<sub>v</sub> | 18 | 6.77 | 2.25E-06 | 2.25E-06 | 9.36 |
|  |  | s-LN<sub>v</sub> | 18 | 12.68 | 9.52E-06 | 9.52E-06 | 11.36 |
|  | LD12:12 | DN<sub>1</sub> | 18 | 27.11 | 4.34E-12 | 4.34E-12 | 60.88 |
|  |  | LN<sub>d</sub> | 21 | 34.74 | 9.60E-17 | 9.60E-17 | 61.2 |
|  |  | I-LN<sub>v</sub> | 19.5 | 21.29 | 2.92E-12 | 2.92E-12 | 38.46 |
|  |  | s-LN<sub>v</sub> | 19.5 | 25.85 | 3.70E-12 | 3.70E-12 | 42 |
|  | LD20:4 | DN<sub>1</sub> | 3 | 10.05 | 0.407078 | 0.407078 | 93.12 |
|  |  | LN<sub>d</sub> | 1.5 | 23.66 | 7.49E-06 | 7.49E-06 | 84.38 |
|  |  | I-LN<sub>v</sub> | 1.5 | 16.62 | 0.296221 | 0.296221 | 71.02 |
|  |  | s-LN<sub>v</sub> | 3 | 15.84 | 7.17E-06 | 7.17E-06 | 38.44 |
| Batumi | LD8:16 | DN<sub>1</sub> | 13.5 | 3.07 | 9.95E-05 | 9.95E-05 | 9.26 |
|  |  | LN<sub>d</sub> | 16.5 | 1.35 | 0.100855 | 0.100855 | 11.25 |
|  |  | I-LN<sub>v</sub> | 16.5 | 0.97 | 0.425168 | 0.425168 | 8.05 |
|  |  | s-LN<sub>v</sub> | 19.5 | 2.79 | 4.44E-07 | 4.44E-07 | 3.35 |
|  | LD12:12 | DN<sub>1</sub> | 16.5 | 1.02 | 0.0089 | 0.0089 | 7.4 |
|  |  | LN<sub>d</sub> | 1.5 | 1.13 | 0.002041 | 0.002041 | 10.53 |
|  |  | I-LN<sub>v</sub> | 1.5 | 1.28 | 0.003109 | 0.003109 | 8.53 |
|  |  | s-LN<sub>v</sub> | 22.5 | 2.17 | 4.10E-06 | 4.10E-06 | 4.3 |
|  | LD20:4 | DN<sub>1</sub> | 18 | 4.11 | 0.825404 | 0.825404 | 32.23 |
|  |  | LN<sub>d</sub> | 21 | 4.86 | 1 | 1 | 37.74 |
|  |  | I-LN<sub>v</sub> | 18 | 2.68 | 1 | 1 | 22.52 |
|  |  | s-LN<sub>v</sub> | 19.5 | 1.17 | 0.263472 | 0.263472 | 3.55 |
| Kilpisjärvi | LD8:16 | DN<sub>1</sub> | 7.5 | 1.51 | 0.797265 | 0.797265 | 9.36 |
|  |  | LN<sub>d</sub> | 12 | 1.83 | 1 | 1 | 17.22 |
|  |  | I-LN<sub>v</sub> | 12 | 2.2 | 0.756486 | 0.756486 | 12.67 |
|  | LD12:12 | DN<sub>1</sub> | 4.5 | 1.11 | 0.304333 | 0.304333 | 11.68 |
|  |  | LN<sub>d</sub> | 16.5 | 1.43 | 0.104619 | 0.104619 | 17.82 |
|  |  | I-LN<sub>v</sub> | 15 | 1.29 | 1 | 1 | 17.76 |
|  | LD20:4 | DN<sub>1</sub> | 12 | 3.41 | 0.589693 | 0.589693 | 29.86 |
|  |  | LN<sub>d</sub> | 16.5 | 8.14 | 0.733528 | 0.733528 | 49.49 |
|  |  | I-LN<sub>v</sub> | 21 | 4.11 | 0.811238 | 0.811238 | 43.16 |

**Table S5.** PDP1 cycling, JTK_CYCLE analysis over a 12h period.

| Strain | Photoperiod | Cluster | Acrophase | Amplitude | P value | Q value | Mean |
| --- | --- | --- | --- | --- | --- | --- | --- |
| *D. melanogaster* | LD8:16 | DN<sub>1</sub> | 3 | 7.111924 | 1 | 1 | 16.41 |
|  |  | LN<sub>d</sub> | 4.5 | 4.52843 | 0.639589 | 0.64 | 19.90 |
|  |  | I-LN<sub>v</sub> | 3 | 3.259085 | 0.18293 | 0.18 | 9.36 |
|  |  | s-LN<sub>v</sub> | 4.5 | 2.451156 | 0.246902 | 0.25 | 11.35 |
|  | LD12:12 | DN<sub>1</sub> | 10.5 | 3.864477 | 0.193058 | 0.19 | 60.87 |
|  |  | LN<sub>d</sub> | 10.5 | 4.549394 | 0.494599 | 0.49 | 61.20 |
|  |  | I-LN<sub>v</sub> | 7.5 | 4.774314 | 0.208045 | 0.21 | 38.45 |
|  |  | s-LN<sub>v</sub> | 10.5 | 3.257501 | 0.027415 | 0.03 | 42 |
|  | LD20:4 | DN<sub>1</sub> | 1.5 | 18.28025 | 0.000195 | 0 | 93.11 |
|  |  | LN<sub>d</sub> | 1.5 | 16.44114 | 0.002375 | 0 | 84.37 |
|  |  | I-LN<sub>v</sub> | 0 | 11.47436 | 0.140583 | 0.14 | 71.01 |
|  |  | s-LN<sub>v</sub> | 1.5 | 8.066222 | 0.018891 | 0.02 | 38.43 |
| Batumi | LD8:16 | DN<sub>1</sub> | 4.5 | 1.570466 | 0.015891 | 0.02 | 9.26 |
|  |  | LN<sub>d</sub> | 4.5 | 1.425012 | 0.358692 | 0.36 | 11.24 |
|  |  | I-LN<sub>v</sub> | 4.5 | 1.453154 | 0.522448 | 0.52 | 8.05 |
|  |  | s-LN<sub>v</sub> | 7.5 | 0.462277 | 0.140322 | 0.14 | 3.34 |
|  | LD12:12 | DN<sub>1</sub> | 1.5 | 0.590659 | 1 | 1 | 7.39 |
|  |  | LN<sub>d</sub> | 7.5 | 0.275419 | 0.778243 | 0.78 | 10.52 |
|  |  | I-LN<sub>v</sub> | NA | 0 | 1 | 1 | 8.53 |
|  |  | s-LN<sub>v</sub> | 0 | 0.917895 | 0.002088 | 0 | 4.29 |
|  | LD20:4 | DN<sub>1</sub> | 3 | 2.869979 | 1 | 1 | 32.23 |
|  |  | LN<sub>d</sub> | 4.5 | 7.639199 | 0.017713 | 0.02 | 37.73 |
|  |  | I-LN<sub>v</sub> | 4.5 | 5.0976 | 0.059576 | 0.06 | 22.52 |
|  |  | s-LN<sub>v</sub> | 3 | 0.971231 | 0.324964 | 0.32 | 3.55 |
| Kilpisjärvi | LD8:16 | DN<sub>1</sub> | 1.5 | 0.89025 | 0.738527 | 0.74 | 9.36 |
|  |  | LN<sub>d</sub> | 9 | 2.014241 | 0.1006 | 0.1 | 17.21 |
|  |  | I-LN<sub>v</sub> | 9 | 2.717112 | 0.157289 | 0.16 | 12.66 |
|  | LD12:12 | DN<sub>1</sub> | 6 | 0.795776 | 1 | 1 | 11.67 |
|  |  | LN<sub>d</sub> | 7.5 | 1.833174 | 0.037607 | 0.04 | 17.81 |
|  |  | I-LN<sub>v</sub> | 7.5 | 0.743767 | 0.992313 | 0.99 | 17.75 |
|  | LD20:4 | DN<sub>1</sub> | 1.5 | 4.996499 | 0.113924 | 0.11 | 29.86 |
|  |  | LN<sub>d</sub> | 3 | 17.33777 | 8.04E-06 | 0 | 49.48 |
|  |  | I-LN<sub>v</sub> | 3 | 14.21752 | 1.69E-06 | 0 | 43.16 |

**Table S6.** PDF DD JTK_CYCLE analysis in the s-LN_v_-terminals.

| Strain | Photoperiod | Acrophase | Amplitude | P value | Q value | Mean |
| --- | --- | --- | --- | --- | --- | --- |
| *D. littoralis*<br>Batumi | DD | 63 | 14.24 | 0.018 | 0.018 | 82.03 |
| *D. littoralis*<br>Raattama | DD | 63 | 3.81 | 1 | 1 | 55.21 |

**Table S7.**
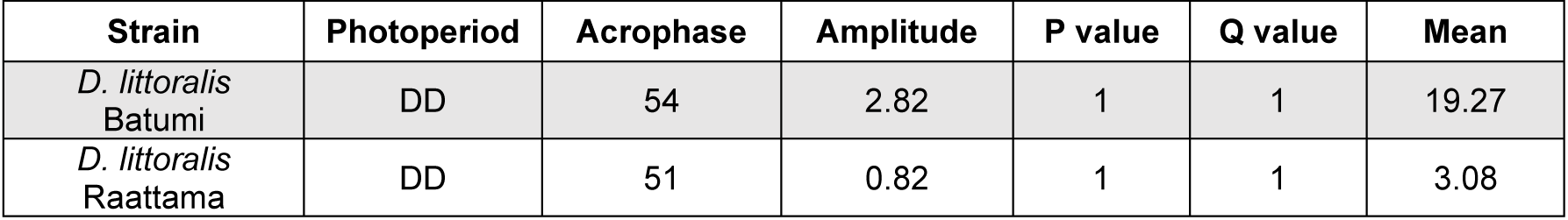
PDP1 DD JTK_CYCLE analysis in the s-LN_v_-nuclei.

| Strain | Photoperiod | Acrophase | Amplitude | P value | Q value | Mean |
| --- | --- | --- | --- | --- | --- | --- |
| *D. littoralis*<br>Batumi | DD | 54 | 2.82 | 1 | 1 | 19.27 |
| *D. littoralis*<br>Raattama | DD | 51 | 0.82 | 1 | 1 | 3.08 |

**Table S8.** JTK cycle analysis for PDF cycling in the s-LN_v_ terminals.

| Strain | Photoperiod | Acrophase | Amplitude | P value | Q value | Mean |
| --- | --- | --- | --- | --- | --- | --- |
| *D. littoralis*<br>Batumi | LD12:12 | 22.5 | 5.82 | 0.006 | 0.006 | 23.32 |

